# Visuospatial attention exerts opposite modulatory effects on explicit and implicit audiovisual subjective synchrony via the frontoparietal network

**DOI:** 10.1101/2025.10.12.681856

**Authors:** Zeliang Jiang, Yuqing Zhao, Weicong Ren, Jie Zhang, Xinyuan Liu, Fei Yu, Xingwei An, Wenxiao Zhong, Pan Zhang, Zhijie Zhang

## Abstract

Synchronization and integration constitute two essential subprocesses of audiovisual binding, commonly indexed by explicit subjective synchrony (ESS) and implicit subjective synchrony (ISS), respectively. Previous studies have shown that spontaneous fluctuations in visual attention are associated with ESS, and that ESS and ISS are negatively correlated, consistent with the temporal renormalization theory. However, it remains unknown whether controlled manipulations of visuospatial attention can causally modulate ESS and ISS in opposite directions via distinct neural mechanisms. To address this question, we conducted two complementary EEG experiments. In Experiment 1, ESS was assessed using a spatially cued beep-flash synchrony judgment task and the point of subjective simultaneity (PSS). In Experiment 2, ISS was measured using fusion and fission variants of the sound-induced flash illusion, with the peak illusion rate (PIR) as the behavioral index. This dual-paradigm design enabled direct comparison between explicit and implicit synchrony under matched attentional manipulations. Behaviorally, asymmetric allocation of visuospatial attention between auditory- and visual-leading conditions induced opposite shifts in PSS and PIR, specifically in the fusion variant. At the neural level, prestimulus frontal theta oscillations were associated with variations in PSS, whereas bilateral frontotemporal and left posterior beta oscillations were associated with variations in PIR. These results demonstrate that visuospatial attention exerts opposite modulatory effects on ESS and ISS through distinct oscillatory dynamics, providing causal evidence for the temporal renormalization theory and advancing understanding of how attentional control shapes multisensory temporal binding.

## 1 Introduction

In noisy environments, successful communication often depends on the ability to simultaneously attend to both visual cues, such as lip movements, and auditory speech signals that are masked by background noise. Such multisensory integration allows listeners to perceive speech more accurately, enhancing message comprehension despite degraded auditory input. The audiovisual integration process can be divided into two distinct subprocesses: synchronization and integration . The synchronization process aligns the temporal aspects of visual and auditory signals and reflects explicit subjective synchrony (ESS), which is typically assessed through conscious judgments of simultaneity. In contrast, the integration process merges visual and auditory inputs into a coherent percept of a single event, reflecting implicit subjective synchrony (ISS), which is typically indexed by automatic, unconscious responses such as reaction times or task accuracy .

With respect to ESS, although some studies suggest that it is an inherent bias shaped by early sensory experiences, multiple studies have shown that ESS can be modulated by more recent perceptual experiences (temporal recalibration effect) and modality-based attention (prior entry effect) . For example, Zampini et al. (2005) quantified ESS using the point of subjective simultaneity (PSS) in an audiovisual synchrony judgment task and found that PSS significantly differed when attention was directed to the visual versus the auditory modality . The underlying logic is that information from the attended modality is prioritized for conscious awareness over information from the unattended modality. This finding supports the view that information from the attended modality gains priority in conscious awareness over that from the unattended modality . Specifically, when attention is directed toward the visual modality, the PSS shifts towards the auditory modality, and conversely, when attention is directed toward the auditory modality, the PSS shifts towards the visual modality.

In addition to modality-based attention, spatial attention allows us to prioritize and selectively process information at specific locations in space . To the best of our knowledge, only one prior study has investigated the effect of endogenous spatial attention on audiovisual temporal integration. In Experiment 3 of that study, a spatially cued audiovisual simultaneity judgment task was employed using simple checkerboard-tone stimuli, with three SOA levels: visual-leading by 300 ms, 150 ms, and 0 ms. The results showed no significant difference in the proportion of simultaneity judgments between attended and unattended conditions . This finding may imply that spatial attention has a limited effect on ESS in the context of simple audiovisual stimuli. A closer examination of the experimental design reveals that the study included only visual-leading SOA conditions, with relatively coarse sampling of SOA levels. As such, it remains unclear whether spatial attention influences simultaneity judgments under auditory-leading SOAs, or under more narrowly spaced SOAs within the typical temporal binding window (TBW; ∼ 200 ms) . A recent study found that, compared to a no-alerting condition, visual alerting significantly improved simultaneity judgment accuracy for auditory-leading stimuli in 120 ms SOAs, as well as for visual-leading stimuli within 100 ms SOAs . Moreover, several studies have demonstrated partially overlapping neural activations between alerting and spatial attention . These findings raise the possibility that visuospatial attention may also modulate audiovisual simultaneity judgments, and thereby influence the ESS. Additional supporting evidence comes from studies on spontaneous attention indexed by prestimulus alpha oscillations . Our study, along with other researchers, has consistently found that lower pre-stimulus spontaneous alpha power is associated with higher synchrony judgment accuracy at the threshold SOAs (the SOAs corresponding to a 50% synchrony judgment rate about ∼100−200 ms). Furthermore, our study revealed a significant positive correlation between the alpha power difference (visual-leading vs. auditory-leading conditions) at threshold SOAs and individuals’ PSS . This finding suggests that the alpha power difference reflects an asymmetry in attentional engagement between the two conditions. A larger alpha difference indicates lower attentional allocation under the visual-leading condition than auditory-leading one, which may reduce synchrony judgment accuracy in that condition. Consequently, the PSS shifts toward the visual-leading side, resulting in a larger PSS value. This finding also suggests a possible effect of spatial attention on ESS.

Unlike in simultaneity judgment tasks, Experiment 1 of Donohue et al. (2015) employed the stream-bounce illusion paradigm and found that, compared to the unattended condition, the proportion of illusory percepts was significantly reduced under spatial attention at visual-leading SOAs of 150 and 300 ms, suggesting that spatial attention inhibit false audiovisual integration, thereby reducing susceptibility to the illusion . In the stream-bounce illusion paradigm, the visual stimuli possess motion characteristics and a natural causal relationship with the accompanying sound. Whether spatial attention exerts a similar modulatory effect on the sound-induced flash illusion (SIFI) with several SOAs— based on simple, static beep-flash stimuli—remains to be further investigated. A review of previous studies reveals that endogenous attention directed to the visual modality has been shown to reduce the occurrence of the fission illusion, while leaving the fusion illusion unaffected . Moreover, another line of research demonstrated that transient disruption of the right posterior parietal cortex — an area implicated in visuospatial attention — increases the likelihood of experiencing the fission illusion . Building on these previous findings, it is plausible that visuospatial attention may influence the sound-induced flash illusion (SIFI), exerting differential effects on the fission and fusion variants and further ISS estimated by illusion rates across different SOAs.

Although ESS plays an important role in audiovisual integration, evidence from both healthy individuals and clinical populations suggests that it is not a sufficient condition for implicit audiovisual integration. Notably, the aforementioned studies employed McGurk illusion stimuli, which contain semantic content; this raises the possibility that higher-level semantic factors may have masked the influence of low-level temporal structure on audiovisual integration . A recent study employing semantically neutral stream-bounce illusion stimuli revealed that the audiovisual SOAs corresponding to ESS and ISS were not aligned. By eliminating semantic influences, this finding provides further evidence that ESS and ISS may rely on distinct underlying processing mechanisms . Interestingly, another study using the same paradigm found a significant negative correlation between ESS and ISS, indicating that although the two are functionally related, they vary in opposite directions . The authors proposed the temporal renormalization theory to account for this negative correlation. This theory posits that the brain contains multiple internal clocks, each of which is biased; however, the average across these clocks provides a relatively accurate estimate of time. This theory highlights that visuospatial attention may exert opposing effects on ESS and ISS.

The deployment of visuospatial attention is regulated by the frontoparietal attention network and is rhythmically modulated through oscillatory activity particularly within the theta, alpha and beta frequency bands. Moreover, neural oscillations at distinct frequencies work in concert to support the temporal and spatial dynamics of multisensory integration . Specifically, theta, alpha and beta oscillations play critical roles in beep-flash based synchrony judgment tasks. While both alpha and beta bands influence the fission variant of the illusion, the fusion variant appears to be more strongly modulated by beta activity . Importantly, attention and audiovisual integration interact at multiple processing stages . Thus, visuospatial attention may influence both ESS and ISS by modulating audiovisual temporal integration through theta to beta oscillatory activity in the frontoparietal network.

In summary, the present study seeks to examine how visuospatial attention modulates both ESS and ISS, and to elucidate the neural oscillatory mechanisms underlying these effects. We conducted two EEG experiments. In Experiment 1, a visuospatially cued audiovisual synchrony judgment task using simple beep-flash stimuli was employed to assess the effect of spatial attention on ESS. In Experiment 2, a visuospatially cued sound-induced flash illusion (SIFI) paradigm — also based on beep-flash stimuli and incorporating both fission and fusion illusion variants — was used to evaluate the influence of spatial attention on ISS. Importantly, we further investigated how oscillatory dynamics prior to stimulus onset — particularly in the theta, alpha and beta frequency bands — contribute to attentional modulation of audiovisual integration across both explicit and implicit levels.

## 2 Material and methods

### 2.1 Participants

Thirty-nine healthy participants (23 females, one left-handed, Mean ± SD age: 21.6 ± 3.2 years) took part in the experiment 1. We initially planned to have the same participants complete both Experiment 1 and Experiment 2. However, due to scheduling conflicts—specifically, some participants from Experiment 1 were unavailable for Experiment 2 due to off-campus internships—we had to recruit additional participants. As a result, Experiment 2 included a total of 36 participants (20 females, all right-handed, Mean ± SD age: 21.3 ± 2.7 years), comprising 16 individuals from Experiment 1 and 20 newly recruited participants. All participants had normal or corrected-to-normal vision and hearing, and reported no history of neurological or psychiatric disorders. The study protocol was approved by the Ethics Committee of Hebei Normal University (approval number: LLSC2025050), and written informed consent was obtained from each participant prior to data collection.

### 2.2 Stimuli

Visual stimuli consisted of white disks (3.4° in diameter, duration=17 ms) presented on a 23.8-inch HP P24V G4 monitor (refresh rate: 60 Hz; resolution: 1920 × 1080). The auditory stimulus was a pure 1800 Hz tone lasting 17 ms (including 3 ms fade-in and fade-out), delivered binaurally through EDIFIER H230P in-ear monitors at a comfortable volume (see Fig. 1).

**Fig. 1.**
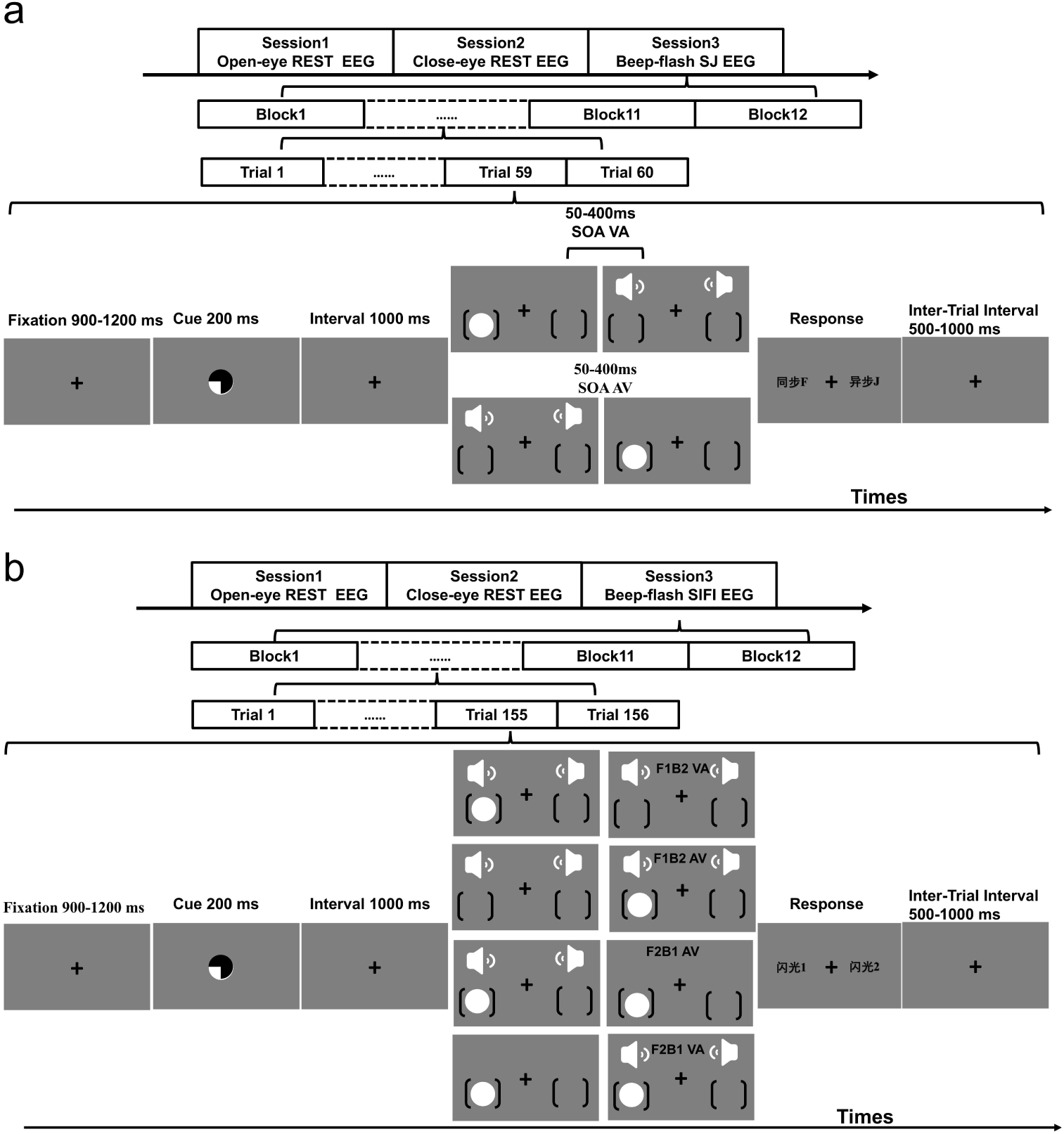
Schematic illustration of the experimental timeline (top) and the sequence of events within a single trial (bottom). (a) for SJ task, while (b) for SIFI task. “AV” indicates visual-leading trials, while “VA” indicates auditory-leading trials. SOAs ranged from 50 to 400 ms (i.e., 50, 100, 200, 300, and 400 ms for SJ task; 50, 100, 200, and 400 ms for SIFI task). F1B2 indicates one flash with two beeps, while F2B1 indicates two flashes with one beep.

### 2.3 Experimental design and procedure

The Experiment 1 used a fully within-subjects factorial design with four factors: (1) cue location (left vs. right), (2) cue validity (valid; attended vs. invalid; unattended), (3) modality order (visual-leading vs. auditory-leading), and (4) SOA (50, 100, 200, 300, or 400 ms). This resulted in a 2×2×2×5 design with 40 total conditions. Each modality order included five SOA levels, yielding 10 distinct audiovisual SOA timing conditions. By convention, SOAs with visual-leading stimuli were coded as positive values, whereas those with auditory-leading stimuli were coded as negative values. Participants completed 720 trials in total, divided into 12 blocks of 60 trials each. Cue validity was manipulated with a 2:1 ratio of valid to invalid trials. Accordingly, each SOA included 48 valid-cue trials and 24 invalid-cue trials. One participant completed an additional block (780 total trials).

The Experiment 2 employed a more complex within-subjects design adapted from previous studies . Each trial presented either one or two visual flashes, paired with zero, one, or two auditory beeps, resulting in six distinct stimulus conditions: (1) one flash with two beeps (F1B2), (2) one flash with one beep (F1B1), (3) one flash with no beep (F1B0), (4) two flashes with two beeps (F2B2), (5) two flashes with one beep (F2B1), and (6) two flashes with no beep (F2B0).

SOA manipulations were applied only to selected conditions. For the fission (F1B2) and fusion (F2B1) illusion conditions, eight SOA values were used: −400, −200, −100, −50, 50, 100, 200, and 400 ms. Negative SOAs indicate auditory-leading sequences, while positive SOAs indicate visual-leading ones. In the F1B2 condition, a negative SOA means the first beep was followed— after a delay— by the flash and second beep; a positive SOA indicates that the first beep and flash were presented together, followed by the second beep. In the F2B1 condition, a negative SOA indicates the first flash and the beep were presented simultaneously, followed by the second flash; a positive SOA means the first flash was presented alone, followed— after a delay— by the second flash and the beep. The F2B0 condition included only four SOA levels (50, 100, 200, and 400 ms), while the F2B2 condition was presented at a fixed SOA of 50 ms. The F1B0 and F1B1 conditions were not temporally manipulated and thus did not include SOA variations.

Experiment 2 primarily focused on the F1B2 and F2B1 conditions, which shared the same design as in Experiment 1, except that the 300 ms SOA was omitted. Participants completed 12 blocks of 156 trials each, for a total of 1872 trials. Within each block, cue location (left vs. right) was balanced (1:1 ratio), and cue validity was manipulated at a 2:1 ratio (valid vs. invalid). Each SOA level appeared in 6 trials per block, while the F1B0 condition appeared in 24 trials, and the F1B1 condition in 6 trials per block.

The overall temporal structure of the experiment was largely consistent between Experiment 1 and Experiment 2. Prior to the main task, 3 minutes of eyes-open resting-state EEG were recorded, followed by 3 minutes of eyes-closed resting-state EEG. Task-related EEG data were then collected subsequently (see Fig. 1). It is worth noting that the resting-state EEG data (eyes-closed and eyes-open) were not included in the subsequent analyses. The task ran on a HP desktop computer with the Psychophysics Toolbox 3.0.11 and MATLAB R2017a (MathWorks).

The experimental procedure of Experiment 1 is described as follows (Fig. 1a). Each trial began with a central fixation cross presented for a jittered duration of 900- 1200 ms [0.51° degree of visual angle (dva)]. This was followed by a symbolic circular cue (1.53° dva) pointing randomly to either the lower left or lower right visual field, presented for 200 ms. After cue offset, there was a 1000 ms interval before stimulus onset. Subsequently, a pair of audiovisual stimuli with varying SOAs was randomly presented. The visual stimulus (3.39° dva) appeared in either the left or right visual field, centered 10.69° dva from the central fixation, while the auditory stimulus was presented binaurally through headphones. Immediately following the second stimulus, a response screen appeared. Participants performed a two-alternative forced-choice simultaneity judgment task, indicating whether the stimuli were synchronous (“F” key) or asynchronous (“J” key). The key-response mapping was counterbalanced across participants. After each response, a blank screen was presented for 500–1000 ms before the onset of the next trial.

The procedural structure of Experiment 1 and Experiment 2 was largely consistent (Fig. 1a). The key difference was that in Experiment 2, participants were tasked with reporting the number of visual flashes (one or two).

### 2.4 EEG recording and preprocessing

The electroencephalogram (EEG) acquisition and preprocessing steps were consistent between Experiment 1 and Experiment 2. The EEG was recorded at 1000 Hz using a actiCHamp amplifier system (Brain Products GmbH, Munich, Germany) with 64 Ag–AgCl scalp electrodes positioned according to the standard international 10–20 system. The online reference was set to the TP9 electrode located near the left mastoid. The ground electrode was positioned between the AF3 and AF4 electrodes. The electrode impedances were maintained below 5 kΩ throughout the recording.

All preprocessing steps were performed using EEGLAB version 13.6.5b . The task-related EEG data were first downsampled to 200 Hz and then filtered using a zero-phase bandpass filter ranging from 0.5 to 80 Hz to remove low- and high-frequency artifacts. A notch filter was subsequently applied to eliminate 50 Hz line noise. The filtered data were re-referenced to the to the average of the left and right mastoid electrodes. The independent component analysis (ICA) was conducted to identify and remove components associated with horizontal and vertical eye movements, as well as muscle artifacts. Finally, EEG data were segmented into epochs ranging from 2 seconds before to 1.5 seconds after the onset of the first stimulus, based on the event markers. Epochs with mean amplitudes exceeding ±100 μV were excluded from further analysis, resulting in an average rejection rate of 5.0 % of trials across participants for Experiment 1 and 1.0 % for Experiment 2.

### 2.5 Behavioral analysis

As a first step, we combined left- and right-cue trials according to cue validity, grouping them into attended (valid cue) and unattended (invalid cue) conditions. Subsequently, we calculated the proportion of synchrony judgments at each SOA in Experiment 1, and the proportions of fission and fusion illusions in Experiment 2, as well as the mean reaction times (RTs) for attended and unattended conditions at each SOA. Subsequently, based on prior studies, we applied MATLAB’s *glmfit* function to fit sigmoid curves and estimate either the PSS or the peak illusion rate (PIR), which were used to quantify ESS and ISS, respectively . The fitting procedure was as follows: First, we applied two separate sigmoid functions to the behavioral response rates for auditory-leading (AV) and visual-leading (VA) trials. The intersection point of these two curves was defined as the PSS or the PIR. To account for non-zero PSS or PIR values, we then re-centered the original SOA data around the estimated PSS or PIR, splitting the data into auditory-leading and visual-leading segments accordingly. The sigmoid fitting was repeated iteratively, updating the split based on the newly estimated intersection point in each iteration, until the two curves converged. Notably, for the fusion illusion, the behavioral response rate at the −400 ms SOA was substantially elevated compared to the −200 ms condition (Fig. 3a), deviating from the expected pattern. This anomaly has been observed in previous studies and was replicated in our own data. Therefore, the −400 ms condition was excluded from the sigmoid fitting to ensure reliable estimation.

Following the curve fitting procedure, each participant obtained four PSS or PIR estimates, corresponding to the following attention conditions: (1) AV attended / VA attended (both attended), (2) AV unattended / VA unattended (both unattended), (3) AV attended / VA unattended, and (4) AV unattended / VA attended. To verify the adequacy of the sigmoid fits, we evaluated the goodness-of-fit for both curves within each of the four experimental conditions. Participants were excluded from further analyses if the goodness-of-fit for either curve in any condition was below 50%. As a result, 11 participants were excluded in Experiment 1 due to poor model fits, resulting in a final sample of 28 participants with an average goodness-of-fit of 91.03 ± 0.08% (mean ± SD). In Experiment 2, 11 participants were excluded under the fusion illusion condition (final N = 25, mean goodness-of-fit = 90.66 ± 5.74%), and 25 participants were excluded under the fission illusion condition (final N = 11, mean goodness-of-fit = 90.67 ± 5.30%).

### 2.6 EEG analysis

All subsequent analyses of the epoched EEG data were conducted using the MATLAB-based Fieldtrip toolbox . Consistent with prior research indicating that contralateral suppression of oscillatory power is a robust indicator of endogenous visuospatial attention, we firstly assessed cue validity in our data by extracting raw EEG signals from regions ipsilateral and contralateral to the cued hemifield. Data from left- and right-cue trials were then collapsed by aligning contralateral and ipsilateral sites across hemispheres. These signals were subjected to time–frequency analysis to determine whether the lateralized neural signature of attentional orienting was reliably elicited.

In previous studies, two primary approaches have been used to analyze EEG data in the context of visuospatial cueing. The most common method involves collapsing left- and right-cue trials into attended and unattended conditions, and examining differences in EEG activity across the whole scalp between these two attentional states . Alternatively, some studies have adopted a lateralized analysis approach, based on the notion that the hemisphere contralateral to the target location has a processing advantage. This method compares EEG activity in the hemisphere contralateral to the target under attended versus unattended conditions . To minimize methodological bias and ensure a comprehensive assessment of attentional effects, we employed both of the above-described approaches in our EEG data analysis. This dual approach allowed us to examine both global and lateralized neural signatures of visuospatial attention.

Time-frequency decomposition was performed using wavelet convolution with the ft_freqanalysis.m function in Fieldtrip. Frequencies ranged from 1 to 30 Hz in 1 Hz increments, with the number of wavelet cycles increasing linearly from 1 cycle (at 1 Hz) to 9 cycles (at 30 Hz). The analysis window extended from −2 to 1.5 s relative to first stimulus (i.e., target) onset. Baseline correction was applied using a reference window from −1.6 to −1.3 seconds relative to target onset (i.e., 300 to 100 ms before cue onset), following protocols from previous studies . Following baseline correction, data were represented as db with respect to the baseline period.

### 2.7 Statistical analysis

#### 2.7.1 Behavioral data statistical analysis

To validate the effectiveness of the attentional manipulation, we first conducted a paired-sample t-test to determine whether RTs were significantly shorter in the attended compared to the unattended condition in both Experiment 1 and Experiment 2. Subsequently, we conducted a 2 (Attention: attended vs. unattended) × 10 (SOA: −400 to 400 ms) repeated-measures ANOVA to investigate whether the proportion of synchrony judgments varied as a function of SOA and attentional state in Experiment 1. A 2 (Attention: attended vs. unattended) × 8 (SOA: − 400 to 400 ms) repeated-measures ANOVA to investigate whether the proportion of illusions varied as a function of SOA and attentional state in Experiment 2. Finally, to test for attentional effects on temporal perception, a one-way repeated-measures ANOVA was performed on PSS or PIR values across four attentional configurations: both attended, both unattended, AV attended / VA unattended, and AV unattended / VA attended. When the assumption of sphericity was violated, the degrees of freedom were corrected using the Greenhouse–Geisser method. Unless otherwise specified, all post hoc comparisons were corrected using Tukey’s Honestly Significant Difference (HSD) method.

#### 2.7.2 EEG statistical analysis

To assess the effectiveness of the attentional cueing, we performed a nonparametric cluster-based permutation test on the time–frequency–channel data. This analysis was implemented using FieldTrip’s ft_freqstatistics function combined with the ft_statfun_depsamplesT method, which applies a paired-sample t-test framework. Multiple comparisons were controlled via a cluster-level correction procedure. The analysis focused on the time window from –1.2 to –0.2 s relative to target onset, capturing cue-induced oscillatory activity while avoiding contamination from early target-evoked responses . Oscillatory power within the 3–30 Hz frequency range was compared between electrode sites contralateral and ipsilateral to the cued location. To unify data across left- and right-cue trials, electrode labels in right-cue trials were flipped so that all right-hemisphere electrodes corresponded to contralateral sites and all left-hemisphere electrodes to ipsilateral sites for left- and right-cue trials. Subsequently, ipsilateral electrode labels were renamed to match their contralateral counterparts, enabling all statistical analyses to be performed within a standardized set of right-hemisphere electrode labels. This approach allowed a direct and consistent comparison of contralateral versus ipsilateral activity regardless of cue direction.

Based on our previous findings that asymmetries in spontaneous attention levels between VA and AV conditions are significantly associated with PSS, we applied a nonparametric cluster-based permutation test on the time–frequency–channel data to further investigate this relationship. This analysis was conducted using FieldTrip’s ft_freqstatistics function combined with the ft_statfun_depsamplesFunivariate statistical method. We compared the power differences between VA and AV conditions in the pre-target interval across four attentional configurations: both attended, both unattended, AV attended / VA unattended, and AV unattended / VA attended, to determine whether these differences were statistically significant. It is worth noting that for this analysis, we focused on a time window from −0.7 to −0.2 s relative to stimulus onset and a frequency range of 3−30 Hz, capturing preparatory oscillatory dynamics prior to stimulus processing.

The nonparametric permutation test proceeded as follows: First, a paired-sample t-test (or repeated-measures ANOVA, depending on the comparison) was performed at each time–frequency–electrode point. Data points exceeding a predefined threshold (*p* < 0.05) were then grouped into clusters based on spatial, temporal, and frequency adjacency, forming candidate clusters of contiguous effects. A cluster was formed only when at least two adjacent points in time, frequency, and electrode space showed significant effects. Next, condition labels were randomly permuted 1000 times to generate a reference distribution of cluster statistics under the null hypothesis. For each permutation, the largest cluster-level statistic was recorded, resulting in an empirical distribution of 1000 values. Finally, a cluster from the observed data was considered significant if its statistic exceeded the 97.5th percentile of the null distribution for t-tests (or the 95th percentile for F-tests), thus correcting for multiple comparisons at the cluster level.

To further explore the neural correlates of behavioral changes, Pearson correlation analyses were conducted between condition-specific PSS or PIR values and oscillatory power metrics that showed significant modulation.

## 3 Results

### 3.1 Behavior Results

#### 3.1.1 Experiment 1

A paired-sample t-test revealed that the RTs were significantly shorter in the attended condition compared to the unattended condition (700.123 ± 29.367 vs. 849.090 ± 45.799 ms; *t*_(38)_ = −5.354, *p* = 4.368 × 10⁻^6^, Cohen’s d = −0.857).

The repeated-measures ANOVA with factors Attention (attended vs. unattended) and SOA (−400 to 400 ms) showed a significant main effect of SOA (*F*_(2.668,101.376)_= 114.370, *p* = 1.600 × 10⁻^97^, partial η² = 0.751), no main effect of Attention (*F*_(1,38)_= 0.327, *p* = 0.571, partial η² = 0.009), but a significant interaction between Attention and SOA (*F*_(5.172,196.528)_= 3.776, *p* = 0.002, partial η² = 0.09). Post-hoc comparisons of the significant Attention × SOA interaction revealed that at the –400 ms SOA, the proportion of synchrony responses was significantly lower in the attended condition compared to the unattended condition (*t*_(38)_ = −2.041, *p* = 0.048, Cohen’s d = −0.327; Fig. 2a). In contrast, at −100, −50, and 50 ms SOAs, the attended condition elicited significantly higher synchrony responses (*t*_(38)_ = 3.008, *p* = 0.005, Cohen’s d = 0.482; *t*_(38)_ = 2.727, *p* = 0.01, Cohen’s d = 0.437; *t*_(38)_ = 2.965, *p* = 0.005, Cohen’s d = 0.475; Fig. 2a). A marginally significant increase was also observed at the 100 ms SOA (*t*_(38)_ = 1.918, *p* = 0.063, Cohen’s d = 0.307; Fig. 2a). No significant differences between attention conditions were found at the remaining SOA levels (all *ps* > 0.1).

**Fig. 2.**
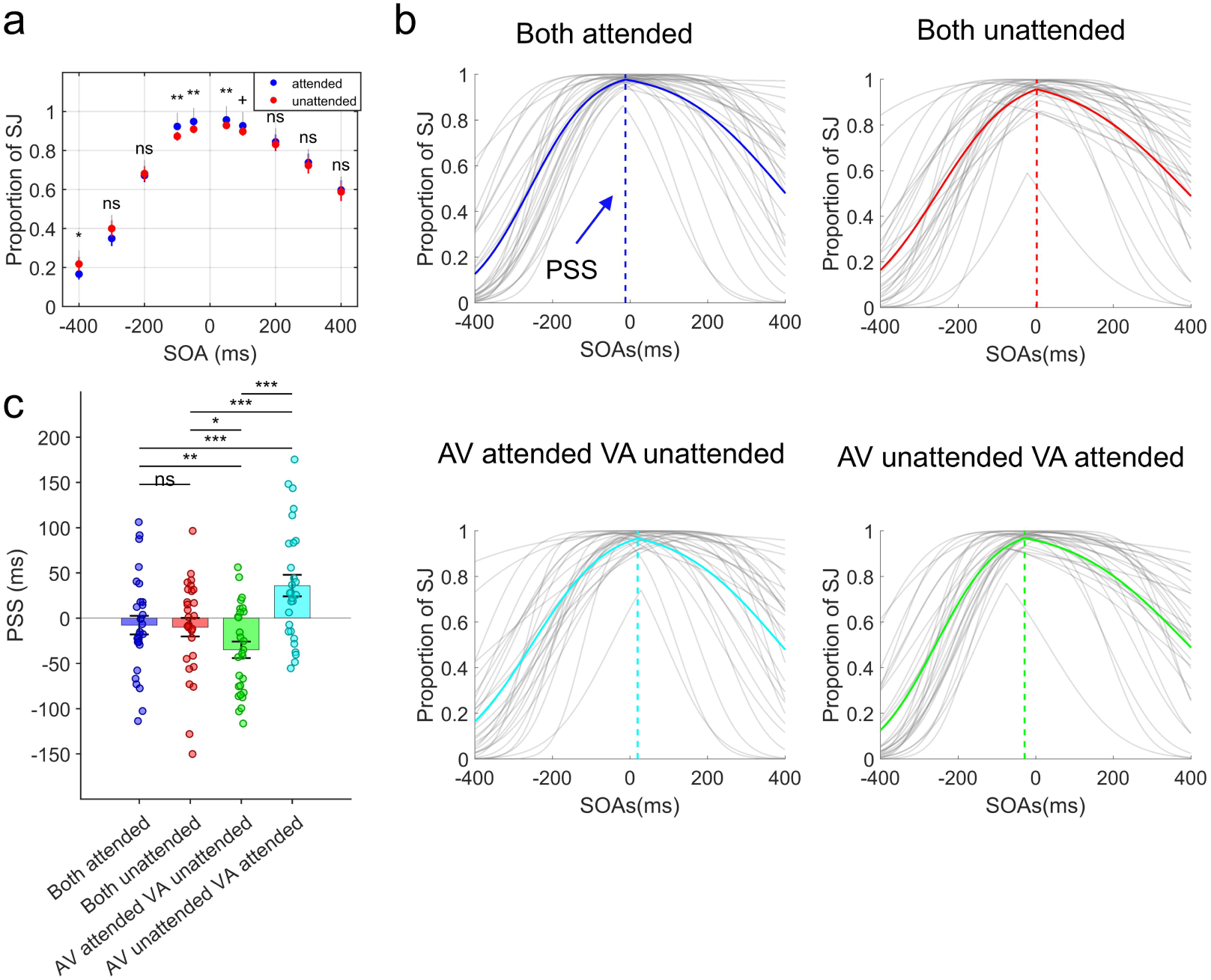
The results of behavioral responses in Experiment 1. (a) The proportion of synchrony judgments across SOAs for attended and unattended conditions, averaged across all participants (N=39). (b) Psychometric curves fitted with dual-sigmoid functions for synchrony responses under four attentional configurations, each shown in a different color (averaged across participants; N=28): blue = both attended, red = both unattended, green = AV attended with VA unattended, cyan = AV unattended with VA attended. Gray lines represent individual participants’ fits. The intersection point between the dashed vertical line and the x-axis indicates the point of subjective simultaneity (PSS). (c) Bar graph showing the mean PSS for each attentional configuration condition, with individual participants represented by dots. Error bars indicate the standard error of the mean. * *p* < 0.05,\*\**p* < 0.001,*** *p* < 0.001,+ *p* < 0.06. ns indicates not significant.

Furthermore, a repeated-measures ANOVA revealed a significant main effect of attentional configuration on the PSS (*F*_(2.300,62.089)_= 13.742, *p* = 4 × 10⁻^6^, partial η² = 0.337; Fig. 2c). Post-hoc comparisons (uncorrected) showed no significant difference in PSS between the both attended and both unattended conditions (*t*_(27)_ = 0.174, *p* = 0.863, Cohen’s d = 0.0329; Fig. 2c). However, PSS values in both of these conditions were significantly larger than those in the AV attended / VA unattended condition (*t*_(27)_ = 3.401, *p* = 0.002, Cohen’s d = 0.643; *t*_(27)_ = 2.687, *p* = 0.012, Cohen’s d = 0.508; Fig. 2c), and significantly smaller than those in the AV unattended / VA attended condition (*t*_(27)_ = −3.780, *p* = 7.890 × 10⁻^4^, Cohen’s d = −0.714; *t*_(27)_ = −3.924, *p* = 5.405 × 10⁻_4_, Cohen’s d = −0.742; Fig. 2c).

#### 3.1.2 Experiment 2

A paired-sample t-test revealed that the RTs were significantly shorter in the attended condition compared to the unattended condition (684.245 ± 22.545 vs. 724.476 ± 25.882 ms, *t*_(35)_ = −5.291, *p* = 6.645 × 10⁻^6^, Cohen’s d = −0.882).

A two-way repeated-measures ANOVA on the raw proportion of fusion illusions revealed significant main effects of both Attention (*F*_(1,35)_= 15.422, *p* = 3.85 × 10⁻^4^, partial η² = 0.306) and SOA (*F*_(2.288,80.097)_= 121.834, *p* = 4.462 × 10⁻^76^, partial η² = 0.777), as well as a significant Attention × SOA interaction (*F*_(3.665,128.281)_= 4.255, *p* = 0.004, partial η² = 0.108; Fig. 3a). Simple effects analysis of the significant Attention × SOA interaction revealed that the proportion of fusion illusions was significantly lower in the attended condition compared to the unattended condition at –200, –100, – 50, 100, and 400 ms SOAs (*t*_(35)_ = −2.867, *p* = 0.007, Cohen’s d = −0.478; *t*_(35)_ = −3.323, *p* = 0.002, Cohen’s d = −0.554; *t*_(35)_ = −2.704, *p* = 0.011, Cohen’s d = −0.451;*t*_(35)_ = −3.293, *p* = 0.002, Cohen’s d = −0.549; *t*_(35)_ = −2.102, *p* = 0.043, Cohen’s d = −0.350; Fig. 3a). In contrast, no significant differences were observed between attention conditions at −400, 50, and 200 ms SOAs (*t*_(35)_ = −0.321, *p* = 0.750, Cohen’s d = −0.054; *t*_(35)_ = −1.612, *p* = 0.116, Cohen’s d = −0.269; *t*_(35)_ = −1.128, *p* = 0.267, Cohen’s d = −0.188; Fig. 3a).

**Fig. 3.**
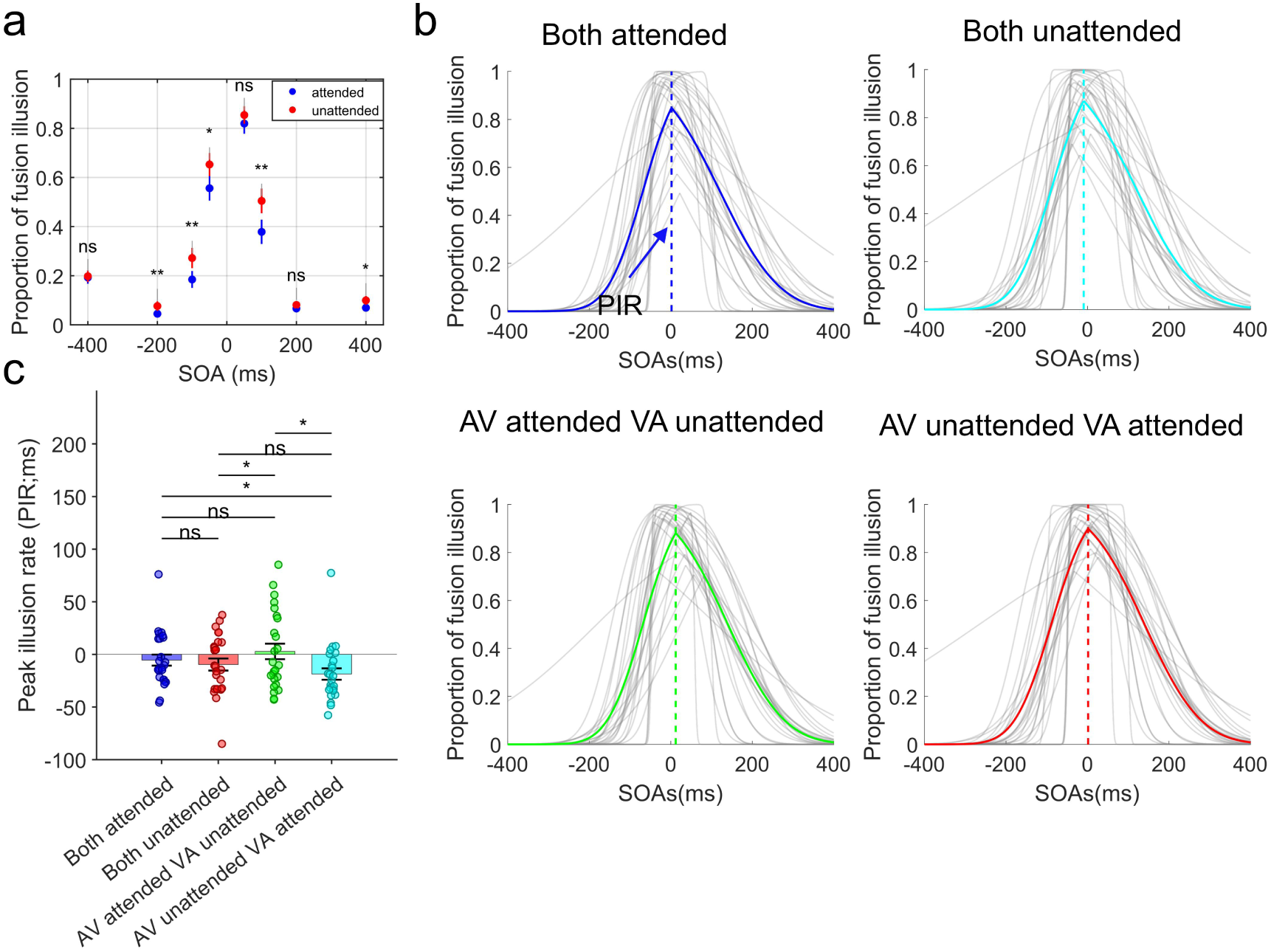
The results of behavioral responses of fusion illusion variant in Experiment 2. (a) The proportion of fusion illusion across SOAs for attended and unattended conditions, averaged across all participants (N=36). (b) Psychometric curves fitted with dual-sigmoid functions for fusion illusion responses under four attentional configurations, each shown in a different color (averaged across participants; N=25): blue = both attended, red = both unattended, green = AV attended with VA unattended, cyan = AV unattended with VA attended. Gray lines represent individual participants’ fits. The intersection point between the dashed vertical line and the x-axis indicates the peak illusion rate (PIR). (c) Bar graph showing the mean PIR for each attentional configuration condition, with individual participants represented by dots. Error bars indicate the standard error of the mean. * *p* < 0.05,\*\**p* < 0.001. ns indicates not significant.

Under the fusion illusion condition, the main effect of attentional configuration on the PIR was significant (*F*_(1.627,39.048)_= 4.501, *p* = 0.024, partial η² = 0.108; Fig. 3c). Post hoc comparisons (uncorrected) revealed that PIR in the both attended condition was significantly higher than in the AV unattended with VA attended condition (*t*_(24)_ = 2.756, *p* = 0.011, Cohen’s d = 0.821; Fig. 3c), while PIR in the both unattended condition was significantly lower than in the AV attended with VA unattended condition (*t*_(24)_ = −2.032, *p* = 0.021, Cohen’s d = −0.466; Fig. 3c). Additionally, PIR in the AV attended with VA unattended condition was significantly higher than in the AV unattended with VA attended condition (*t*_(24)_ = 3.183, *p* = 0.023, Cohen’s d = 0.458; Fig. 3c). No other pairwise comparisons reached statistical significance (all *ps* > 0.1; Fig. 3c).

Under the fission illusion condition, a two-way repeated-measures ANOVA on the raw illusion rates revealed significant main effects of both attention (*F*_(1,35)_= 4.384, *p* = 0.044, partial η² = 0.111) and SOA (*F*_(2.229,78.023)_= 17.957, *p* = 3.350 × 10⁻^19^, partial η² = 0.339). However, the interaction between attention and SOA was not significant (*F*_(4.710,164.837)_= 0.595, *p* = 0.694, partial η² = 0.017; Fig. 4a). Post hoc comparisons for the attention factor revealed that the proportion of fission illusions was significantly lower in the attended condition compared to the unattended condition (*t*_(35)_ = −2.094, *p* = 0.044, Cohen’s d = -0.349). Under the fission illusion condition, the main effect of attentional configuration on PIR was significant (*F*_(1.627,39.048)_= 4.501, *p* = 0.024, partial η² = 0.108; Fig. 4c). Post hoc comparisons (uncorrected) revealed that PIR in the both attended condition was significantly higher than in the AV unattended with VA attended condition (*t*_(10)_ = 2.658, *p* = 0.024, Cohen’s d = 0.502; Fig. 4c), and that PIR in the AV attended with VA unattended condition was also significantly higher than in the AV unattended with VA attended condition (*t*_(10)_ = 2.633, *p* = 0.025, Cohen’s d = 0.498; Fig. 4c). No other pairwise differences reached significance (all *ps* > 0.1).

**Fig. 4.**
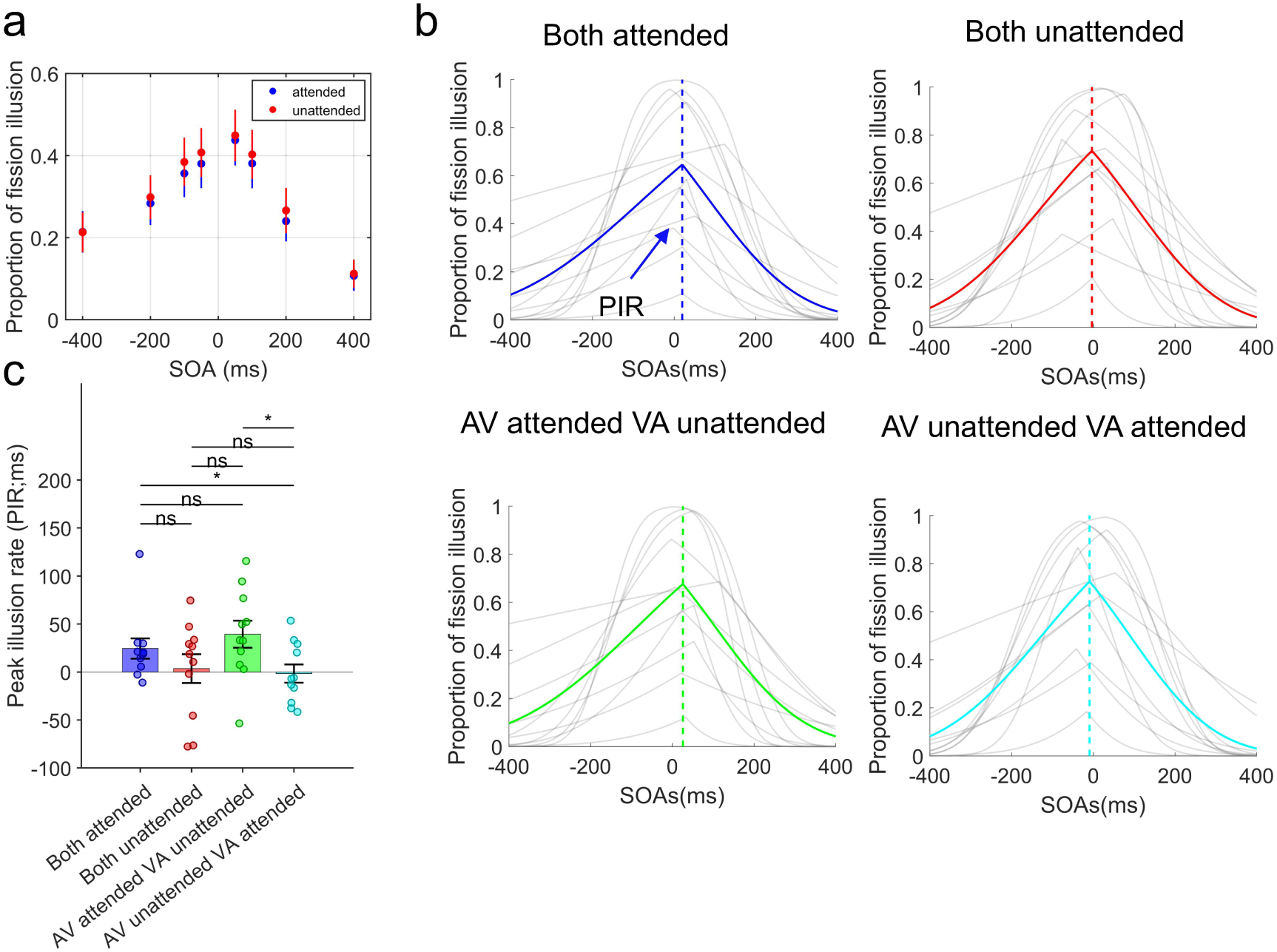
The results of behavioral responses of fission illusion variant in Experiment 2. (a) The proportion of fission illusion across SOAs for attended and unattended conditions, averaged across all participants (N=36). (b) Psychometric curves fitted with dual-sigmoid functions for fission illusion responses under four attentional configurations, each shown in a different color (averaged across participants; N=11): blue = both attended, red = both unattended, green = AV attended with VA unattended, cyan = AV unattended with VA attended. Gray lines represent individual participants’ fits. The intersection point between the dashed vertical line and the x-axis indicates the peak illusion rate (PIR). (c) Bar graph showing the mean PIR for each attentional configuration condition, with individual participants represented by dots. Error bars indicate the standard error of the mean. * *p* < 0.05. ns indicates not significant.

### 3.2 EEG results

#### 3.2.1 Experiment 1

To validate cue effectiveness, we conducted a nonparametric permutation test using paired-sample t-tests on time–frequency power from contralateral and ipsilateral electrodes following cue onset. The analysis identified a significant negative cluster predominantly distributed over central-parietal, parietal-occipital, frontal, and mid- frontal electrode sites (cluster value = −4.947 × 10_4_, *p* = 0.002; Fig. S1a). In the central-parietal and parietal-occipital regions, this cluster was mainly observed between −0.9 and −0.4 s in the 5–16 Hz frequency range (Fig. S1b). In the frontal and mid-frontal regions, it was primarily distributed between −0.8 and −0.2 s in the 20–30 Hz range (Fig. S1b).

To investigate the neural oscillatory power associated with visuospatial attention that contributes to variations in PSS across different attentional configuration conditions, we conducted a one-way ANOVA across all scalp electrodes using a non- parametric cluster-based permutation test. This analysis revealed a significant positive cluster (cluster value = 1.244 × 10_4_, *p* = 0.029). This cluster was primarily distributed over the frontal, frontocentral, mid-parietal, and parietal electrodes. In the frontal and frontocentral regions (AFz, AF3, AF4, AF7, AF8, F1, F2, F3, F4, F6, F7, FCz, FC1, FC2, FC3, FC4, FC5, FC6; Fig. 5a), the significant effects were observed in the 5–11 Hz frequency range during the −0.6 to −0.55 s time window. In contrast, in the mid- parietal and parietal regions (CP1, CP3, CP4, CP5, CP6, P1, P3, P4, P5, P6, P8; Fig. 5c), the effects were evident in the 3–7 Hz frequency range within the same time window.

**Fig. 5.**
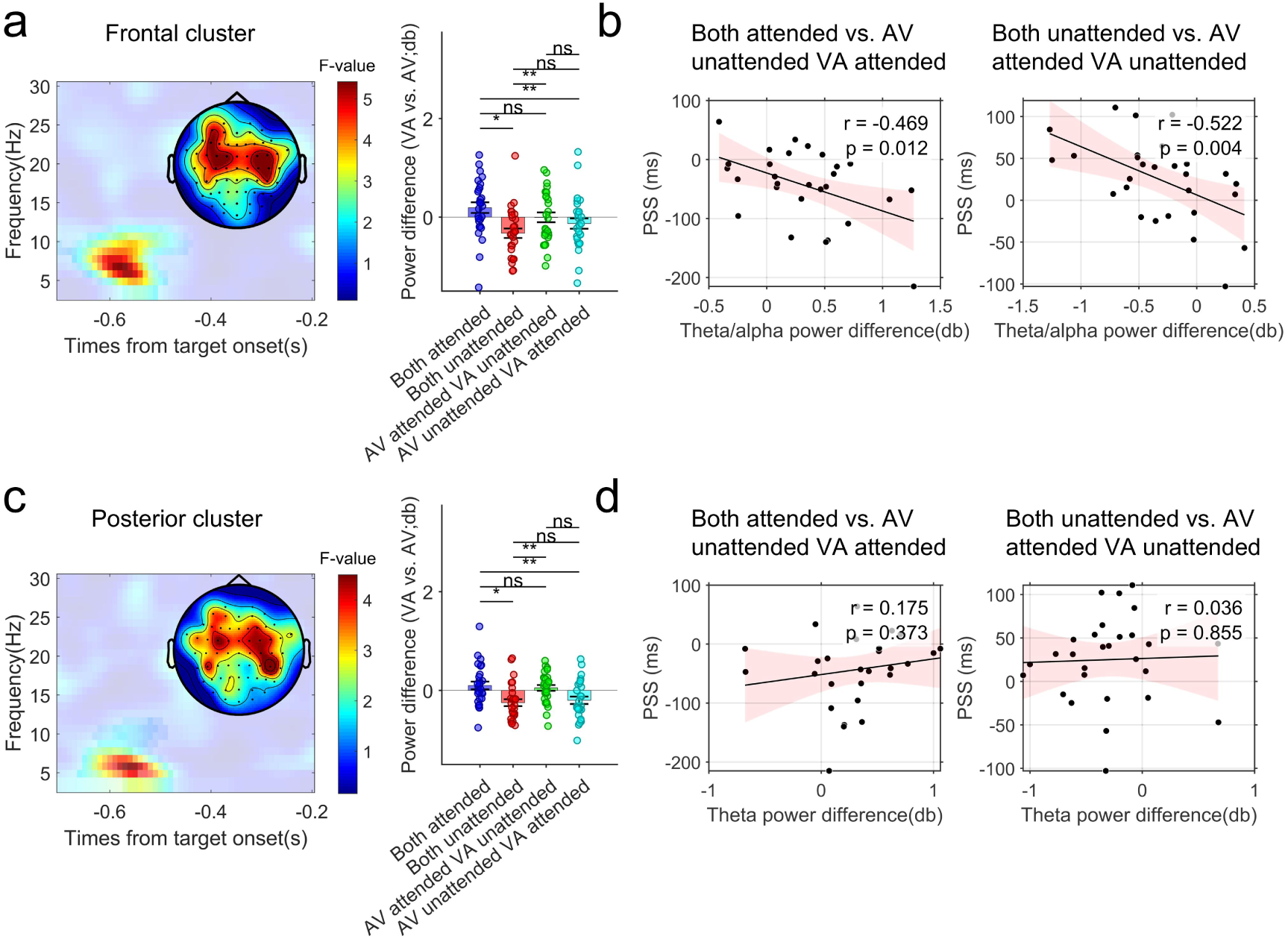
Prestimulus Oscillatory Power Differences Across Attentional Configurations and Their Relationship with Behavioral Performance in the whole-brain analysis. (a) and (c) illustrate the main effects of attentional configuration on pre-stimulus oscillatory power at frontal ([AFz, AF3, AF4, AF7, AF8, F1, F2, F3, F4, F6, F7, FCz, FC1, FC2, FC3, FC4, FC5, FC6]) and posterior ([CP1, CP3, CP4, CP5, CP6, P1, P3, P4, P5, P6, P8]) electrodes, respectively, as represented by F-value time-frequency plots. Unmasked regions indicate significant time– frequency clusters; masked areas represent non-significant results. The accompanying bar graphs show the results of post-hoc comparisons, illustrating how oscillatory power varied across different attentional configuration levels. The topographical map in (a) shows the scalp distribution of F-values averaged over the 5–11 Hz frequency range and –0.60 to –0.55 s time window. The corresponding map in (b) displays the F-value distribution averaged over the 3–7 Hz frequency range and the same time window. (b) and (d) show scatter plots of Pearson correlations between the differences in oscillatory power (for the significantly different levels revealed in the post-hoc tests in (a) and (c)) and the corresponding differences in PSS. Each black dot represents an individual participant, the solid line depicts the linear regression fit, and the light pink shaded area indicates the 95% confidence interval of the fit. * *p* < 0.05,\*\**p* < 0.01. ns indicates not significant.

We averaged oscillatory power difference (VA vs. AV) within the identified time–frequency windows and conducted post hoc simple effects analyses (HSD- corrected). In the frontal and frontocentral electrodes, the VA–AV power difference was significantly larger in the both attended condition compared to the both unattended condition (*t*_(27)_ = 3.198, *p* = 0.017, Cohen’s d = 0.604; Fig. 5a) and the AV unattended VA attended condition (*t*_(27)_ = 3.843, *p* = 0.004, Cohen’s d = 0.726; Fig. 5a). Additionally, the both unattended condition showed a significantly smaller VA– AV power difference than the AV attended / VA unattended condition (*t*_(27)_ = -3.843, *p* = 0.004, Cohen’s d = –0.726; Fig. 5a). No other pairwise comparisons reached significance (all *ps* > 0.1). Given the significant PSS differences between the both attended and AV unattended / VA attended conditions, as well as between the both unattended and AV attended / VA unattended conditions, we conducted Pearson correlation analyses between the neural power differences and the corresponding PSS differences in these two contrasts. The results revealed significant negative correlations in both cases (both attended vs. AV unattended VA attended: *r*_(26)_ = −0.469, *p*=0.012; both unattended vs. AV attended VA unattended: *r*_(26)_ = −0.522, *p*=0.004; Fig. 5b). Notably, the power differences in these contrasts corresponded to the difference in oscillatory power between the attended and unattended conditions specifically under the AV condition. Therefore, these correlation results suggest that, larger power in the AV unattended compared to the AV attended condition is associated with increased PSS, indicating a stronger bias toward the VA direction, whereas smaller power in the AV attended compared to the AV unattended condition is associated with decreased PSS, indicating a stronger bias toward the AV direction.

In the mid-parietal and parietal electrodes, the VA–AV power difference was also significantly larger in the both attended condition compared to the both unattended condition (*t*_(27)_ = 2.877, *p* = 0.037, Cohen’s d = 0.544; Fig. 5c) and the AV unattended / VA attended condition (*t*_(27)_ = 3.948, *p* = 0.003, Cohen’s d = 0.746; Fig. 5c). Additionally, the both unattended condition showed a significantly smaller VA– AV power difference than the AV attended VA unattended condition (*t*_(27)_ = −3.948, *p* = 0.003, Cohen’s d = −0.746; Fig. 5c). No other pairwise comparisons reached significance (all *ps* > 0.1). However, no significant correlations were observed between the power differences and PSS differences in any of the conditions (both attended vs. AV unattended / VA attended: *r*_(26)_ = 0.175, *p* = 0.373; both unattended vs. AV attended / VA unattended: *r*_(26)_ = 0.036, *p* = 0.855). This suggests that although attention configuration influenced both neural oscillatory power in posterior electrodes and PSS, their relationship was not directly correlated across participants.

To further explore whether the attentional modulation of PSS was lateralized, and in contrast to the whole-brain analysis described above, we conducted a one-way ANOVA restricted to electrodes contralateral to the target using a non-parametric cluster-based permutation test. As a result, we identified two significant positive clusters (cluster1 value = 6.995 × 10_3_, *p* = 0.016; cluster2 value = 5.015 × 10_3_, *p* = 0.045). The first cluster closely resembled that identified in the whole-brain analysis, spanning frontal electrodes (AF4, AF8, F2, F4, F6, FC2, FC4, FC6) from −0.6 to −0.55 s in the 5–11 Hz frequency range (Fig. S2a), and posterior electrodes (CP4, CP6, P4, P6, P8) from −0.7 to −0.55 s in the 3–7 Hz range (Fig. S2c). The second cluster was primarily distributed over parietal and parieto-occipital electrodes (P2, P4, P6, P8, PO4, PO8), spanning from −0.7 to −0.6 s in the 11–19 Hz frequency range (Fig. S2e).

Post-hoc analyses on Cluster 1 in the frontal region revealed that the VA–AV power difference was significantly larger in the both attended condition than in the both unattended condition (*t*_(27)_ = 2.831, *p* = 0.041, Cohen’s d = 0.535; Fig. S2a) and the AV unattended / VA attended condition (*t*_(27)_ = 2.888, *p* = 0.036, Cohen’s d = 0.546). Moreover, the both unattended condition exhibited a significantly smaller VA– AV power difference compared to the AV attended / VA unattended condition (*t*_(27)_ = – 2.888, *p* = 0.036, Cohen’s d = –0.546). No other pairwise comparisons reached statistical significance (all *ps* > 0.4). In line with the whole-brain findings, Pearson correlation analyses further demonstrated a robust negative association between power differences across attentional configuration conditions and changes in PSS (both attended vs. AV unattended / VA attended: *r*_(26)_ = –0.371, *p* = 0.026, one-tailed; both unattended vs. AV attended / VA unattended: *r*_(26)_ = –0.468, *p* = 0.012; Fig. S2b). Post-hoc comparisons of posterior electrodes in Cluster 1 showed the same pattern as in the frontal region: the VA–AV power difference was larger in the both attended condition than in the both unattended condition (*t*_(27)_ = 2.747, *p* = 0.049, Cohen’s d = 0.519; Fig. S2c) and the AV unattended / VA attended condition (*t*_(27)_ = 3.931, *p* = 0.003, Cohen’s d = 0.743; Fig. S2c), and was smaller in the both unattended condition compared to the AV attended / VA unattended condition (*t*_(27)_ = – 3.931, *p* = 0.003, Cohen’s d = -0.743; Fig. S2c). As in the whole-brain analysis, we did not observe any significant correlations between power differences across conditions and corresponding PSS differences (both attended vs. AV unattended VA attended: *r*_(26)_ = -0.082, *p* = 0.677; both unattended vs. AV attended VA unattended: *r*_(26)_ = -0.183, *p* = 0.350; Fig. S2d).

Post-hoc analysis of the raw power differences in the posterior electrodes of Cluster 2 revealed no significant difference between the both attended and both unattended conditions (*t*_(27)_ = 0.358, *p* = 0.984, Cohen’s d = 0.068; Fig. S2e). However, the power difference in the both attended condition was significantly smaller than in the AV attended / VA unattended condition (*t*_(27)_ = –3.168, *p* = 0.019, Cohen’s d = – 0.599; Fig. S2e) and significantly larger than in the AV unattended / VA attended condition (*t*_(27)_ = 4.042, *p* = 0.002, Cohen’s d = 0.764; Fig. S2e). Likewise, the both unattended condition showed a significantly smaller power difference than the AV attended / VA unattended condition (*t*_(27)_ = –4.042, *p* = 0.002, Cohen’s d = –0.764; Fig. S2e). Furthermore, the AV attended / VA unattended condition exhibited a significantly larger power difference compared to the AV unattended / VA attended condition (*t*_(27)_ = 4.500, *p* = 6.390 × 10^-4^, Cohen’s d = 0.850). No significant correlations were observed between the power differences across conditions and the corresponding PSS differences (all *ps* > 0.3), suggesting that the neural responses within this cluster may not underlie the attentional modulation of PSS.

#### 3.2.2 Experiment 2

Consistent with Experiment 1, the non-parametric permutation test revealed that post-cue oscillatory power in the contralateral hemisphere was significantly lower than that in the ipsilateral hemisphere. However, unlike Experiment 1, the significant cluster (cluster value = −1.717 × 10_4_, *p* = 0.002; Fig. S3a) was more spatially confined, primarily distributed over mid-parietal, parietal, and parieto-occipital electrodes within the 9–19 Hz frequency range and the −0.92 to −0.45 s time window (Fig. S3b).

A non-parametric cluster-based permutation test based on a within-subjects one- way ANOVA revealed significant differences in the VA–AV power difference across different attentional configurations (cluster value = 9.490 × 10_3_, *p* = 0.049). This cluster was primarily distributed within the time window of −0.5 to −0.3 s and the frequency range of 17 to 22 Hz. Spatially, it was localized over the bilateral frontal and frontotemporal electrodes (F7, FT7, F8, FT8; Fig. 6a), as well as the left mid- parietal and parieto-occipital electrodes (CP5, CP3, P5, P3, PO3; Fig. 6b). As shown in Figures 6a and 6c, the most significant F-values were observed in the time-frequency window of −0.5 to −0.4 s and 17–22 Hz. Therefore, we averaged the VA– AV power differences across this window for the frontal and posterior parietal electrodes within the significant cluster and conducted post-hoc comparisons across attentional configurations respectively.

**Fig. 6.**
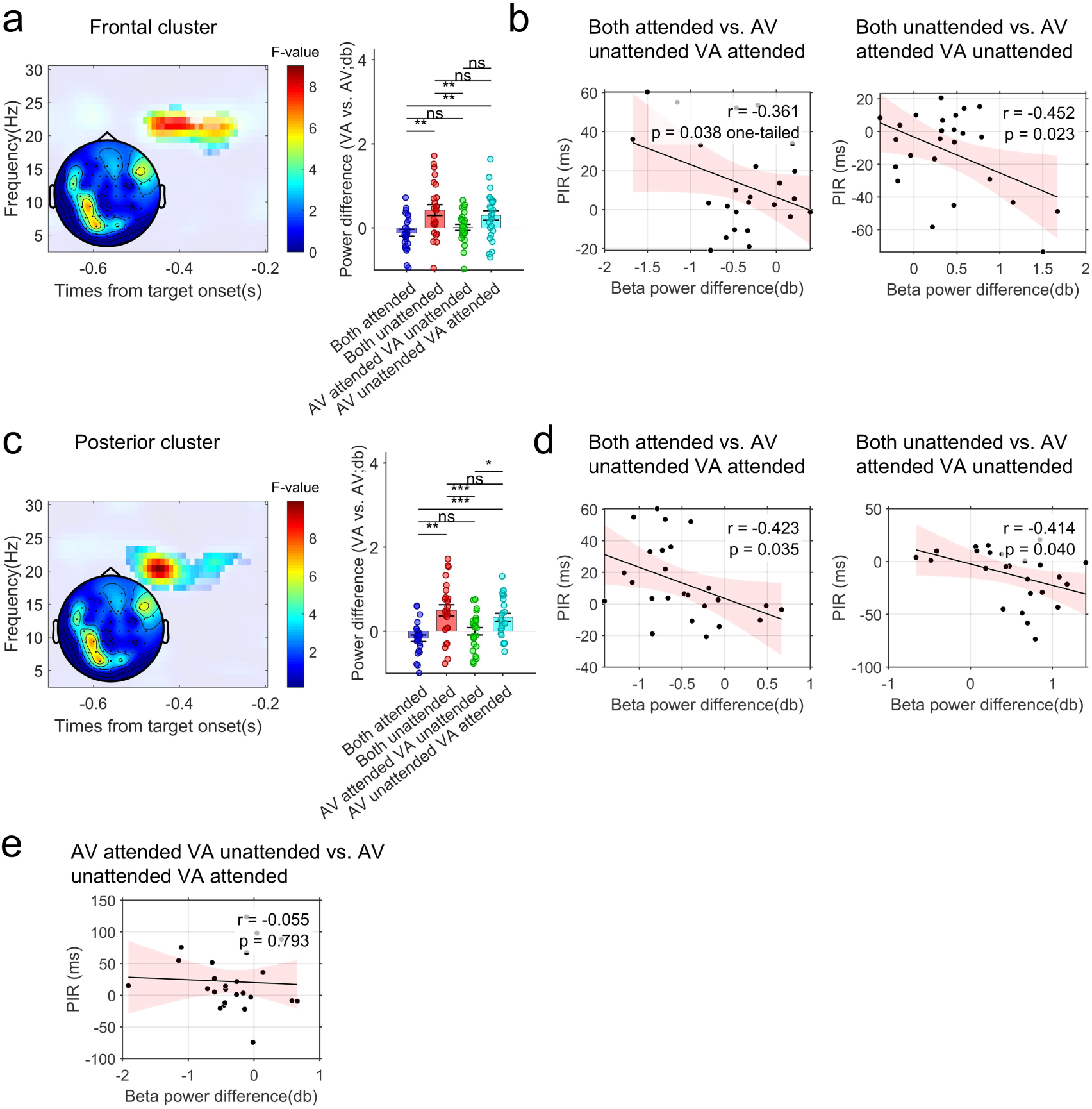
Prestimulus Oscillatory Power Differences Across Attentional Configurations and Their Relationship with Behavioral Performance in the whole-brain analysis. (a) and (c) illustrate the main effects of attentional configuration on pre-stimulus oscillatory power at frontal ([F8, FT8, F7, FT7]) and posterior ([CP5, CP3, P5, P3, PO3]) electrodes, respectively, as represented by F-value time-frequency plots. Unmasked regions indicate significant time– frequency clusters; masked areas represent non-significant results. The accompanying bar graphs show the results of post-hoc comparisons, illustrating how oscillatory power varied across different attentional configuration levels. The topographical map in (a) and (b) shows the scalp distribution of F-values averaged over the 17–22 Hz frequency range and –0.50 to –0.4 s time window. (b), (d), and (e) show scatter plots of Pearson correlations between the differences in oscillatory power (for the significantly different levels revealed in the post-hoc tests in (a) and (c)) and the corresponding differences in PIR. Each black dot represents an individual participant, the solid line depicts the linear regression fit, and the light pink shaded area indicates the 95% confidence interval of the fit. * *p* < 0.05,\*\**p* < 0.01. ns indicates not significant.

Post-hoc analysis of the frontal electrodes revealed that the VA–AV power difference was significantly smaller in the both attended condition compared to the both unattended condition (*t*_(24)_ = –3.493, *p* = 0.010, Cohen’s d = –0.660; Fig. 6a) and the AV unattended / VA attended condition (*t*_(24)_ = –3.989, *p* = 0.003, Cohen’s d = – 0.754; Fig. 6a). Additionally, the power difference in the both unattended condition was significantly larger than that in the AV attended / VA unattended condition (*t*_(24)_ = 3.989, *p* = 0.003, Cohen’s d = 0.754; Fig. 6a). Subsequent correlation analysis showed that larger power differences were significantly associated with smaller PIR differences (both attended vs. AV unattended VA attended: *r*_(23)_ = -0.361, *p* = 0.038, one-tailed; both unattended vs. AV attended VA unattended: *r*_(23)_ = -0.452, *p* = 0.023; Fig. 6b), indicating a negative relationship between the two measures. Consistent with Experiment 1, the power differences observed in the correlation analysis primarily reflect the contrast between the AV attended and unattended conditions. Combined with these correlation results, it can be inferred that, relative to the AV attended condition, decreased power in the AV unattended condition directly leads to a reduction in PIR. Conversely, compared to the AV unattended condition, increased power in the AV attended condition directly results in an increase in PIR.

Post-hoc analysis of the posterior electrodes revealed a pattern similar to that of the frontal sites. The VA–AV power difference was significantly larger in both the both unattended condition (*t*_(24)_ = –3.958, *p* = 0.003, Cohen’s d = –0.748; Fig. 6c) and the AV unattended / VA attended condition (*t*_(24)_ = –4.815, *p* = 3.642 × 10^-4^, Cohen’s d = –0.910; Fig. 6c), compared to the AV attended / VA unattended condition. Moreover, the power difference in the both unattended condition was significantly larger than that in the AV attended / VA unattended condition (*t*_(24)_ = 4.815, *p* = 3.642 × 10_-4_, Cohen’s d = 0.910; Fig. 6c). Additionally, the power difference in the AV attended / VA unattended condition was significantly smaller than in the AV unattended / VA attended condition (*t*_(24)_ = –3.005, *p* = 0.029, Cohen’s d = –0.568; Fig. 6c). Consistent with the correlation results in the frontal region, a significant negative association between power difference and PSS difference was also observed in the posterior region. However, this relationship was only found for the power difference between the both attended and the AV unattended / VA attended conditions (*r*_(23)_ = –0.423, *p* = 0.035; Fig. 6d), as well as between the both unattended and the AV attended / VA unattended conditions (*r*_(23)_ = –0.414, *p* = 0.040; Fig. 6d). No significant correlation was observed between the AV attended / VA unattended and the AV unattended / VA attended conditions (*r*_(23)_ = –0.055, *p* = 0.793; Fig. 6e).

Consistent with Experiment 1, we conducted hemisphere-wise ANOVAs on power differences across attention configurations. A cluster approaching statistical significance was observed over parietal and parieto-occipital electrodes (cluster value = 4.234 × 10_3_, *p* = 0.075).

As shown in Fig. S3, the marginal significance may stem from the lateralized attentional modulation being restricted to posterior electrodes. Including additional electrode sites likely increased the number of comparisons, thereby elevating the risk of false negatives. To address this, we averaged the data from mid-parietal, parietal, and parieto-occipital electrodes (CP4, CP6, TP8, P2, P4, P6, P8, PO4, PO8) and re-ran a one-way ANOVA. The analysis revealed a significant main effect of attention configuration (cluster value = 835.754, *p* = 0.012; Fig. S4a). A prominent concentration of F-values within the cluster was observed between –0.6 to –0.5 s and 23 to 26 Hz. Post hoc tests on the extracted within the time-frequency cluster revealed that the VA–AV power difference was significantly smaller in the both attended condition compared to both the both unattended condition (*t*_(24)_ = –3.792, *p* = 0.005, Cohen’s d = –0.717; Fig. S4a) and the AV unattended / VA attended condition (*t*_(24)_ = –2.970, *p* = 0.031, Cohen’s d = –0.561; Fig. S4a). Furthermore, the power difference in the both unattended condition was significantly larger than that in the AV attended / VA unattended condition (*t*_(24)_ = 2.970, *p* = 0.031, Cohen’s d = 0.561; Fig. S4a). However, Pearson correlation analyses between power differences across attention configurations and PSS differences revealed no significant associations (both attended vs. AV unattended VA attended: *r*_(23)_ = –0.316, *p* = 0.123; both unattended vs. AV attended VA unattended: *r*_(23)_ = –0.318, *p* = 0.121; Fig. S4b).

## 4 Discussion

Synchrony and integration are two essential sub-processes of multisensory integration. Previous studies have revealed multiple interactions between attention and multisensory integration: multisensory integration can enhance bottom-up attention, while top-down attention can, in turn, facilitate multisensory integration. However, no study to date has directly examined the influence of attention on these two sub-processes— synchrony and integration. In the present study, we employed a cued audiovisual SJ task and a SIFI task combined with beep-flash stimuli to investigate whether, and how, top-down visuospatial attention modulates the processes of synchrony and integration. The synchrony process was quantified using the ESS (i.e., PSS) derived from the SJ task, whereas the integration process was indexed by the ISS (i.e., PIR) in the SIFI task. Our behavioral results demonstrate that visuospatial attention exerts opposite effects on ESS and ISS. Specifically, when the levels of attention in the AV and VA conditions were equivalent—either both attended or both unattended—no significant differences were observed in PSS or PIR between the two conditions. In contrast, when attention was directed to the AV condition but not to the VA condition, the PSS was significantly reduced, whereas the PIR was significantly increased. Conversely, when attention was absent in the AV condition but present in the VA condition, the PSS was significantly increased, while the PIR was significantly reduced. The EEG results revealed that low-frequency oscillatory power in the frontal and parietal regions (theta, alpha, and beta bands) reflected the level of attention, thereby directly contributing to the observed changes in PSS and PIR.

### 4.1 Effectiveness of Attentional Cueing Manipulation

In this study, endogenous visual cues were employed to direct participants’ attention to specific visuospatial locations, enabling us to examine behavioral differences between attended and unattended conditions. The effectiveness of the attentional manipulation is a prerequisite for the reliability of the present findings. Following previous research, we assessed the validity of this manipulation by comparing behavioral RTs and EEG oscillatory power between the attended and unattended conditions. The behavioral results showed that, in both Experiment 1 and Experiment 2, RTs in the attended condition were significantly shorter than those in the unattended condition. This finding is consistent with previous studies and indicates that valid cues facilitated faster behavioral responses. The EEG results revealed that, relative to electrodes ipsilateral to the attended location, alpha-band power and power in adjacent frequency bands over posterior electrodes contralateral to the attended location were significantly reduced. This pattern replicates findings reported in multiple previous studies . According to the *gating by inhibition* theory, the increase in alpha power ipsilateral to the cue reflects the brain’s suppression of irrelevant information, whereas the decrease in alpha power contralateral to the cue reflects the disinhibition of relevant information, thereby facilitating its processing. Notably, compared with Experiment 2, Experiment 1 also revealed a significant reduction in theta- and beta-band power over contralateral frontal electrodes (Fig. S1). This finding is not surprising, as previous studies have similarly identified decreases in contralateral frontal theta- and beta-band power as a hallmark of visuospatial attention . Based on the frontal electrode distribution of this effect, we infer that it may originate from the frontal eye fields within the dorsal attention network . A possible reason why this effect was not observed in Experiment 2 is that the SIFI task was relatively simple and required a relatively lower level of attentional investment . In summary, the behavioral and EEG results from both Experiment 1 and Experiment 2 collectively demonstrate that the attentional manipulation employed in this study was effective.

### 4.2 Visuospatial Attention Exerts Opposite Effects on ESS and ISS

In Experiment 1, we found that under 100 ms SOA conditions or shorter, the proportion of synchronous responses in the attended condition was significantly higher than that of asynchronous responses in both auditory-leading and visual-leading conditions, whereas at the auditory-leading 400 ms SOA, the pattern was reversed (Fig. 2a). Donohue et al. (2015) reported that endogenous spatial attention does not affect audiovisual simultaneity judgments for simple stimuli . The present study extends this finding by showing that the influence of visuospatial attention on simultaneity judgments is dependent on SOA duration. Specifically, within the TBW (∼100−200 ms), attention facilitated synchronous judgments, whereas outside this TBW—particularly in auditory-leading SOAs—attention promoted asynchronous judgments. Given that participants tend to integrate audiovisual stimuli within the TBW and to segregate them outside this window, these results suggest that attention can facilitate both audiovisual integration and segregation. This conclusion is consistent with previous findings that visuospatial attention can exert dual influences on temporal processing, facilitating both the integration and segregation of visual events depending on task demands . By fitting the raw synchrony response rates, we obtained the PSS values for different attentional configurations. ANOVA and post-hoc tests revealed no significant difference in PSS between the both attended and both unattended conditions. However, the PSS in these two conditions was significantly larger than in the AV attended / VA unattended condition and smaller than in the AV unattended / VA attended condition. Moreover, the PSS in the AV attended / VA unattended condition was significantly smaller than in the AV unattended / VA attended condition. Although the number of participants included in the fitting-based analysis of PSS (N=28) was smaller than that in the analysis of the raw synchrony response rates (N=39), the PSS results closely mirrored the differences between attended and unattended conditions across SOAs observed in the original proportion data. As shown in Figures 2a and 2b, compared with the both unattended condition, the both attended condition—where attention was present in both AV and VA trials—produced an increase in the absolute slopes of the fitted curves for both AV and VA, with the intersection point shifting upward rather than horizontally. Similarly, in the AV attended / VA unattended condition, relative to the both attended or both unattended conditions, the absolute slope of the VA or AV curve decreased or increased, and the PSS shifted leftward, resulting in a smaller PSS. In the AV unattended / VA attended condition, the pattern was exactly reversed. Several previous studies have defined the slope of the fitted function as an index of temporal sensitivity, and one study even directly regarded it as a measure of the TBW . From this perspective, the present results suggest that attention can enhance temporal sensitivity—or equivalently, narrow the TBW—under both visual-leading and auditory-leading SOA conditions, thereby influencing ESS.

In Experiment 2, under the fusion illusion variant, we found that the raw proportion of fusion illusions was significantly lower in the attended condition than in the unattended condition for multiple SOA conditions, except for the auditory-leading 400 ms SOA and the visual-leading 50 ms and 200 ms SOAs (Fig. 3a). Under the fission illusion variant, when collapsing across all SOA conditions, we observed that the proportion of fission illusions was also significantly lower in the attended condition than in the unattended condition (Fig. 4a). The results of Experiment 1 in Donohue et al. (2015) showed that, in the stream–bounce illusion paradigm, the proportion of “bounce” responses was significantly lower in the attended condition than in the unattended condition at SOAs of 150 ms and 300 ms . Our findings extend this result to the SIFI paradigm, indicating that attention can indeed suppress erroneous audiovisual integration. At first glance, this result may appear inconsistent with Experiment 1, where the attended condition exhibited an elevated synchrony response rate at SOAs within the TBW. However, the two findings are actually consistent: in Experiment 1, attention facilitated correct audiovisual integration within the TBW, whereas in Experiment 2, the decrease in illusion rates under the attended condition indicates that attention suppressed erroneous audiovisual integration. Further analysis of the PIR values obtained from the fitted data revealed that, particularly under the fusion variant, there was no significant difference between the both unattended and both attended conditions. The PIR in the both attended condition was significantly greater than in the AV unattended / VA attended condition, whereas the PIR in the both unattended condition was significantly smaller than in the AV attended / VA unattended condition (Fig. 3c). These results closely mirrored the distribution of the raw illusion proportions under attended and unattended conditions. Moreover, the slope changes observed in the fitted curves can also account for these findings as in the interpretation of the results from Experiment 1 (Fig. 3b and 4b). Given that only 11 participants met the fitting criteria for the PIR in the fission variant, the relatively small sample size limits the statistical power of the analysis. Therefore, future studies with larger samples are needed to determine whether results similar to those observed in the fusion variant can be obtained under the fission variant.

Taken together, the behavioral results from both experiments indicate that visuospatial attention indeed modulates audiovisual integration, although its manifestation differs between explicit and implicit paradigms. In the explicit paradigm, attention facilitates audiovisual integration by increasing synchrony responses within the TBW and suppresses erroneous audiovisual integration by decreasing synchrony responses outside the TBW, thereby promoting audiovisual segregation. In the implicit paradigm, attention suppresses erroneous audiovisual integration by reducing the proportion of illusions across almost all SOA conditions. This pattern is also reflected in the effects on ESS and ISS, particularly in the AV attended / VA unattended and AV unattended / VA attended conditions, where the influence of attention on ESS and ISS is exactly opposite. According to the Temporal Renormalization Theory, both ESS and ISS are subject to biases, and their mean reflects veridical temporal perception . Our findings provide indirect support for this theory.

### 4.3 Oscillatory Power in the Frontoparietal Network Influences both ESS and ISS

In Experiment 1, whole-scalp electrodes analysis revealed that the frontal theta/alpha and the posterior parietal theta power difference (VA−AV) were significantly larger in the both attended condition than in the AV unattended / VA attended condition, and significantly smaller in the both unattended condition than in the AV attended / VA unattended condition (Fig. 5a). Importantly, correlation analysis revealed that the difference in frontal power difference (VA−AV) between the both attended condition and the AV unattended / VA attended condition— specifically, the frontal power difference between AV unattended and AV attended— was significantly negatively correlated with the PSS difference between these two conditions (Fig. 5c). A similar negative correlation was observed between the both unattended condition and the AV attended / VA unattended condition (Fig. 5d). These results directly indicate that higher frontal power under AV unattended conditions is associated with a rightward shift (increase) in PSS, whereas lower frontal power under AV attended trials is associated with a leftward shift (decrease) in PSS. In the analysis of electrodes contralateral to the target location, we replicated the findings observed in the whole-scalp analysis. Given the frontoparietal distribution of the observed power effects, we infer that they are part of the dorsal attention network, encompassing regions such as the frontal eye fields and the intraparietal sulcus . Although we additionally observed differences in posterior alpha power across attentional configurations, correlation analysis did not reveal a direct relationship between these differences and PSS changes. Although numerous previous studies have shown that frontal theta power is closely associated with cognitive control, a recent study reported that, compared with neutral cues, visuospatial attention elicits higher prestimulus frontal theta power, indicating its involvement in spatial attention . In addition, studies have reported that audiovisual bimodal attention or cross-modal selective attention elicits a significant increase in post-stimulus frontal theta power, indicating its involvement in cross-modal attentional processes. In the present study, although we employed visuospatial cues, participants were also required to attend to the auditory stimuli. Therefore, the frontal theta/alpha activity observed here may be related to both visuospatial attention and cross-modal attention. However, it is noteworthy that previous studies have consistently reported that attention is associated with increases in frontal power, whereas in the present study, we observed a decrease in power. One possible explanation is that prior research primarily focused on post-stimulus evoked oscillations, whereas the present study examined pre-stimulus spontaneous oscillations. Post-stimulus evoked power is measured relative to baseline, and a reduction in pre-stimulus spontaneous power effectively lowers this baseline, which can in turn produce an apparent increase in evoked power. In other words, decreases in pre-stimulus power and increases in post- stimulus power may reflect the same underlying neural process. No relationship was observed between power changes and PSS differences in the VA attended versus unattended conditions. This may reflect that attention does not directly modulate behavioral responses in VA condition, but exerts its influence indirectly via post- stimulus neural oscillations.

Unlike in Experiment 1, Whole-brain analysis in Experiment 2 revealed significant differences in beta band power across attention configurations over the bilateral lateral frontal as well as the left middle parietal and parieto-occipital channels. Post-hoc tests showed that the beta band power difference (VA–AV) in the both attended condition was significantly smaller than in the AV unattended / VA attended condition, whereas in the both unattended condition, the beta band power difference was significantly larger than in the AV attended / VA unattended condition. In other words, beta band power in the AV attended condition was significantly higher than in the AV unattended condition. Further correlation analyses revealed that the beta band power difference between the both attended condition and the AV unattended / VA attended condition was significantly negatively correlated with the corresponding difference in PIR. Likewise, the beta band power difference between the both unattended condition and the AV attended / VA unattended condition also exhibited a significant negative correlation with the PIR difference. These findings suggest that beta band power in the AV attended and AV unattended conditions directly modulates changes in PIR. Specifically, smaller beta band power in the AV unattended condition was associated with smaller PIR values, whereas greater beta band power in the AV attended condition was associated with larger PIR values. Notably, analysis of the hemisphere contralateral to the target revealed no significant association between power differences and behavioral differences across attention configurations. This finding suggests that the attentional modulation of PIR does not exhibit a contralateral processing advantage.

In light of previous literature and considering the topographical distribution of the beta band effects, we infer that the effects observed over the bilateral lateral frontal electrodes may originate from the inferior frontal gyrus (IFG), whereas those over the middle parietal and parieto-occipital electrodes may arise from the intraparietal sulcus (IPS). Previous extensive evidence indicates that the IFG constitutes a core component of the ventral attention network involved in bottom-up attention, whereas the IPS serves as a key hub of the dorsal attention network engaged in top-down attention . In this study, visuospatial attention was of a top-down nature, and our focus was primarily on neural activity preceding stimulus onset. Therefore, we suggest that the involvement of the IFG does not reflect a bottom-up attentional process. Recent fMRI studies have indeed demonstrated that the IFG is also involved in top-down attentional processes . It has been proposed that the IFG may function as a relay, transmitting signals from the dorsal attention network (DAN) to the visual cortex, and may also contribute to interpreting visual cues and orienting attention . In research on vision and attention, beta-band activity is considered the predominant neural rhythm of the dorsal visual pathway, particularly within the parietal cortex. It is thought to provide precise spatial coordinates for visual perception, thereby facilitating visual processing . In general, beta-band activity is implicated in top- down processing and is associated with the maintenance of the current cognitive state . Using the same paradigm, Keil et al. (2014) found that beta-band power in the left temporal region was significantly higher for flash-illusion responses than for non-illusion responses. This finding also suggests that pre-stimulus spontaneous beta-band activity is associated with maintaining the cognitive state underlying the illusion. Interestingly, our results revealed that beta-band power increased under the attention condition, while the proportion of illusion responses decreased, suggesting that participants may have utilized elevated beta activity to maintain a non-illusory perceptual state. In summary, the beta-band activity observed in the present study reflects top-down attentional processing, which may provide precise spatial coordinates and support the maintenance of spatial attention. Since the illusion task did not require participants to attend to the auditory stimuli, it is unlikely to have engaged bimodal attention, which may explain the absence of theta-band effects in Experiment 2.

Overall, the EEG findings from the two Experiments suggest that visuospatial attention influences explicit synchrony and implicit illusion judgments—and thereby modulates ESS and ISS—via low-frequency (∼30 Hz) oscillations in the frontoparietal network. The specific frequency bands involved in explicit and implicit tasks depend on the nature of the experimental task.

### 4.4 Limitations and Future Directions

It is important to acknowledge two limitations of this study. First, in Experiment 1, although the maximum audiovisual SOA was extended to 400 ms, participants’ average proportion of simultaneity responses in the visual-leading 400-ms SOA condition remained around 0.6. This suggests that peripheral visual stimuli posed a relatively high perceptual challenge, and that this SOA still fell within participants’ TBW. Consequently, unlike in the auditory-leading condition, we were unable to determine whether simultaneity judgments at SOAs clearly outside the TBW were modulated by visuospatial attention. Second, due to inconsistent goodness-of-fit across attentional configurations, approximately 30 % of participants in the fusion- variant conditions of Experiments 2 and 1 were excluded based on fitting criteria, and this proportion increased to nearly 70% in the fission-variant condition of Experiment 2. As a result, the statistical effect sizes for PSS and PIR were relatively small. Moreover, no post-hoc correction was applied in the behavioral statistical analyses. Therefore, future studies should employ a denser and more systematically distributed sampling of SOAs to validate and strengthen the reliability of the present findings.

## 5 Conclusion

In this study, we employed two EEG experiments to investigate how visuospatial attention modulates explicit (ESS) and implicit (ISS) synchrony perception, as well as the underlying neural oscillatory mechanisms. Behaviorally, we found that when attention was asymmetrically allocated between visual-leading and auditory-leading conditions, ESS and ISS shifted in opposite directions compared with conditions of symmetrical attention allocation. This pattern indirectly supports the temporal renormalization theory. At the neural level, prestimulus oscillatory activity revealed that ESS modulation was associated with theta-band power in frontal electrodes, whereas ISS modulation was linked to beta-band power in bilateral frontotemporal and left mid-posterior electrodes electrodes. Taken together, these findings suggest that visuospatial attention exerts opposite modulations on ESS and ISS through the engagement of the frontoparietal network.

## Acknowledgments

This work was partially supported by the Doctoral Research Start-up Fund Project (No.13102164) and the Digital Humanities Special Project of Hebei Normal University (No. S25SZ003).

## Reference

[1] Freeman ED, Ipser A, Palmbaha A, et al. Sight and sound out of synch: Fragmentation and renormalisation of audiovisual integration and subjective timing[J]. cortex, 2013, 49(10): 2875–2887.

[2] Scurry AN, Vercillo T, Nicholson A, et al. Aging impairs temporal sensitivity, but not perceptual synchrony, across modalities[J]. Multisensory research, 2019, 32(8): 671–692.

[3] Zhou H, Cheung EF, Chan RC. Audiovisual temporal integration: Cognitive processing, neural mechanisms, developmental trajectory and potential interventions[J]. Neuropsychologia, 2020, 140: 107396.

[4] Grabot L, Van Wassenhove V. Time Order as Psychological Bias[J]. Psychological Science, 2017, 28(5): 670–678.

[5] Badde S, Ley P, Rajendran SS, et al. Sensory experience during early sensitive periods shapes cross-modal temporal biases[J]. elife, 2020, 9: e61238.

[6] Fujisaki W, Shimojo S, Kashino M, et al. Recalibration of audiovisual simultaneity[J]. Nature neuroscience, 2004, 7(7): 773–778.

[7] Van der Burg E, Alais D, Cass J. Rapid recalibration to audiovisual asynchrony[J]. Journal of neuroscience, 2013, 33(37): 14633–14637.

[8] Zampini M, Shore DI, Spence C. Audiovisual prior entry[J]. Neuroscience letters, 2005, 381(3): 217–222.

[9] Spence C, Parise C. Prior-entry: A review[J]. Consciousness and cognition, 2010, 19(1): 364– 379.

[10] Titchener EB. Lectures on the elementary psychology of feeling and attention[M]. Macmillan, 1908.

[11] Carrasco M. How visual spatial attention alters perception[J]. Cognitive Processing, 2018, 19(S1): 77–88.

[12] Donohue SE, Green JJ, Woldorff MG. The effects of attention on the temporal integration of multisensory stimuli[J]. Frontiers in integrative neuroscience, 2015, 9: 32.

[13] Zampini M, Guest S, Shore DI, et al. Audio-visual simultaneity judgments[J]. Perception & Psychophysics, 2005, 67(3): 531–544.

[14] Bastiaansen M, Berberyan H, Stekelenburg JJ, et al. Are alpha oscillations instrumental in multisensory synchrony perception?[J]. Brain Research, 2020, 1734: 146744.

[15] Li Q, Liu P, Huang S, et al. The influence of phasic alerting on multisensory temporal precision[J]. Experimental Brain Research, 2018, 236(12): 3279–3296.

[16] Sturm W, Schmenk B, Fimm B, et al. Spatial attention: more than intrinsic alerting?[J]. Experimental Brain Research, 2006, 171(1): 16–25.

[17] Chandrakumar D, Keage HA, Gutteridge D, et al. Interactions between spatial attention and alertness in healthy adults: a meta-analysis[J]. Cortex, 2019, 119: 61–73.

[18] Corbetta M, Shulman GL. Spatial Neglect and Attention Networks[J]. Annual Review of Neuroscience, 2011, 34(1): 569–599.

[19] Jensen O, Mazaheri A. Shaping functional architecture by oscillatory alpha activity: gating by inhibition[J]. Frontiers in human neuroscience, 2010, 4: 186.

[20] Klimesch W. Alpha-band oscillations, attention, and controlled access to stored information[J]. Trends in cognitive sciences, 2012, 16(12): 606–617.

[21] Dehaghani NS, Zarei M. Pre-stimulus activities affect subsequent visual processing: Empirical evidence and potential neural mechanisms[J]. Brain and Behavior, 2025, 15(2).

[22] Jiang Z, An X, Liu S, et al. Beyond alpha band: prestimulus local oscillation and interregional synchrony of the beta band shape the temporal perception of the audiovisual beep-flash stimulus[J]. Journal of Neural Engineering, 2024, 21(3): 036035.

[23] Jiang Z, An X, Liu S, et al. Spontaneous alpha-band oscillations reflect individual differences in audiovisual temporal perception[C]//2023 45th Annual International Conference of the IEEE Engineering in Medicine & Biology Society (EMBC). IEEE, 2023: 1–4.

[24] Vroomen J, Keetels M. Perception of causality and synchrony dissociate in the audiovisual bounce-inducing effect (ABE)[J]. Cognition, 2020, 204: 104340.

[25] Zhang M, Tang X, Yu W, et al. The effects of modal-based endogenous attention on sound-induced flash illusion.[J]. Acta Psychologica Sinica, 2018.

[26] Cattaneo Z, Silvanto J, Pascual-Leone A, et al. The role of the angular gyrus in the modulation of visuospatial attention by the mental number line[J]. Neuroimage, 2009, 44(2): 563–568.

[27] Kamke MR, Vieth HE, Cottrell D, et al. Parietal disruption alters audiovisual binding in the sound-induced flash illusion[J]. NeuroImage, 2012, 62(3): 1334–1341.

[28] Soto-Faraco S, Alsius A. Deconstructing the McGurk–MacDonald illusion.[J]. Journal of Experimental Psychology: Human perception and performance, 2009, 35(2): 580.

[29] Martin B, Giersch A, Huron C, et al. Temporal event structure and timing in schizophrenia: preserved binding in a longer “now”[J]. Neuropsychologia, 2013, 51(2): 358–371.

[30] Tsilionis E, Vatakis A. Multisensory binding: Is the contribution of synchrony and semantic congruency obligatory?[J]. Current Opinion in Behavioral Sciences, 2016, 8: 7–13.

[31] Scolari M, Seidl-Rathkopf KN, Kastner S. Functions of the human frontoparietal attention network: Evidence from neuroimaging[J]. Current opinion in behavioral sciences, 2015, 1: 32–39.

[32] Tosoni A, Capotosto P, Baldassarre A, et al. Neuroimaging evidence supporting a dual- network architecture for the control of visuospatial attention in the human brain: a mini review[J]. Frontiers in Human Neuroscience, 2023, 17: 1250096.

[33] Fiebelkorn IC, Pinsk MA, Kastner S. A dynamic interplay within the frontoparietal network underlies rhythmic spatial attention[J]. Neuron, 2018, 99(4): 842–853.

[34] Gaillard C, Ben Hamed S. The neural bases of spatial attention and perceptual rhythms[J]. European Journal of Neuroscience, 2022, 55(11–12): 3209–3223.

[35] Foster JJ, Sutterer DW, Serences JT, et al. Alpha-Band Oscillations Enable Spatially and Temporally Resolved Tracking of Covert Spatial Attention[J]. Psychological Science, 2017, 28(7): 929–941.

[36] Foster JJ, Awh E. The role of alpha oscillations in spatial attention: limited evidence for a suppression account[J]. Current opinion in psychology, 2019, 29: 34–40.

[37] Yang X, Fiebelkorn IC, Jensen O, et al. Differential neural mechanisms underlie cortical gating of visual spatial attention mediated by alpha-band oscillations[J]. Proceedings of the National Academy of Sciences, 2024, 121(45): e2313304121.

[38] Buschman TJ, Miller EK. Top-Down Versus Bottom-Up Control of Attention in the Prefrontal and Posterior Parietal Cortices[J]. Science, 2007, 315(5820): 1860–1862.

[39] Lundqvist M, Miller EK, Nordmark J, et al. Beta: bursts of cognition[J]. Trends in Cognitive Sciences, 2024, 28(7): 662–676.

[40] Keil J, Senkowski D. Neural Oscillations Orchestrate Multisensory Processing[J]. The Neuroscientist, 2018, 24(6): 609–626.

[41] Bastiaansen MCM, Berberyan H, Stekelenburg J, et al. Neural oscillations in the perception of audiovisual synchrony[C]//NVP Winter Conference. 2017.

[42] Kambe J, Kakimoto Y, Araki O. Phase reset affects auditory-visual simultaneity judgment[J]. Cognitive Neurodynamics, 2015, 9(5): 487–493.

[43] Yuan X, Li H, Liu P, et al. Pre-stimulus beta and gamma oscillatory power predicts perceived audiovisual simultaneity[J]. International Journal of Psychophysiology, 2016, 107: 29–36.

[44] Keil J, Senkowski D. Individual alpha frequency relates to the sound-induced flash illusion[J]. Multisensory Research, 2017, 30(6): 565–578.

[45] Keil J, Müller N, Hartmann T, et al. Prestimulus beta power and phase synchrony influence the sound-induced flash illusion[J]. Cerebral Cortex, 2014, 24(5): 1278–1288.

[46] Noguchi Y. Individual differences in beta frequency correlate with the audio–visual fusion illusion[J]. Psychophysiology, 2022, 59(8): e14041.

[47] Koelewijn T, Bronkhorst A, Theeuwes J. Attention and the multiple stages of multisensory integration: A review of audiovisual studies[J]. Acta psychologica, 2010, 134(3): 372–384.

[48] Talsma D, Senkowski D, Soto-Faraco S, et al. The multifaceted interplay between attention and multisensory integration[J]. Trends in cognitive sciences, 2010, 14(9): 400–410.

[49] Tang X, Wu J, Shen Y. The interactions of multisensory integration with endogenous and exogenous attention[J]. Neuroscience & Biobehavioral Reviews, 2016, 61: 208–224.

[50] Gavin N, Hirst RJ, McGovern DP. The magnitude of the sound-induced flash illusion does not increase monotonically as a function of visual stimulus eccentricity[J]. Attention, Perception, & Psychophysics, 2022, 84(5): 1689–1698.

[51] Delorme A, Makeig S. EEGLAB: an open source toolbox for analysis of single-trial EEG dynamics including independent component analysis[J]. Journal of neuroscience methods, 2004, 134(1): 9–21.

[52] Stevenson RA, Wallace MT. Multisensory temporal integration: task and stimulus dependencies[J]. Experimental Brain Research, 2013, 227(2): 249–261.

[53] Ipser A, Karlinski M, Freeman ED. Correlation of individual differences in audiovisual asynchrony across stimuli and tasks: New constraints on temporal renormalization theory.[J]. Journal of Experimental Psychology: Human Perception and Performance, 2018, 44(8): 1283.

[54] Oostenveld R, Fries P, Maris E, et al. FieldTrip: Open Source Software for Advanced Analysis of MEG, EEG, and Invasive Electrophysiological Data[J]. Computational Intelligence and Neuroscience, 2011, 2011: 1–9.

[55] Van der Lubbe RH, Utzerath C. Lateralized power spectra of the EEG as an index of visuospatial attention[J]. Advances in cognitive psychology, 2013, 9(4): 184.

[56] Busch NA, VanRullen R. Spontaneous EEG oscillations reveal periodic sampling of visual attention[J]. Proceedings of the National Academy of Sciences, 2010, 107(37): 16048– 16053.

[57] Yang X, Ying C, Zhu L, et al. The neural oscillations in delta- and theta-bands contribute to divided attention in audiovisual integration[J]. Perception, 2024, 53(1): 44–60.

[58] Harris AM, Dux PE, Mattingley JB. Detecting unattended stimuli depends on the phase of prestimulus neural oscillations[J]. Journal of Neuroscience, 2018, 38(12): 3092–3101.

[59] Gould IC, Rushworth MF, Nobre AC. Indexing the graded allocation of visuospatial attention using anticipatory alpha oscillations[J]. Journal of Neurophysiology, 2011, 105(3): 1318– 1326.

[60] Gallotto S, Duecker F, Ten Oever S, et al. Relating alpha power modulations to competing visuospatial attention theories[J]. NeuroImage, 2020, 207: 116429.

[61] Grabot L, Kayser C. Alpha activity reflects the magnitude of an individual bias in human perception[J]. Journal of Neuroscience, Society for Neuroscience, 2020, 40(17): 3443–3454.

[62] Thut G, Nietzel A, Brandt SA, et al. α-Band electroencephalographic activity over occipital cortex indexes visuospatial attention bias and predicts visual target detection[J]. Journal of neuroscience, 2006, 26(37): 9494–9502.

[63] Gould IC, Rushworth MF, Nobre AC. Indexing the graded allocation of visuospatial attention using anticipatory alpha oscillations[J]. Journal of Neurophysiology, 2011, 105(3): 1318– 1326.

[64] Wildegger T, Van Ede F, Woolrich M, et al. Preparatory α-band oscillations reflect spatial gating independently of predictions regarding target identity[J]. Journal of Neurophysiology, 2017, 117(3): 1385–1394.

[65] Gallotto S, Duecker F, Ten Oever S, et al. Relating alpha power modulations to competing visuospatial attention theories[J]. NeuroImage, 2020, 207: 116429.

[66] Yang X, Fiebelkorn IC, Jensen O, et al. Differential neural mechanisms underlie cortical gating of visual spatial attention mediated by alpha-band oscillations[J]. Proceedings of the National Academy of Sciences, 2024, 121(45): e2313304121.

[67] Marshall TR, O’Shea J, Jensen O, et al. Frontal eye fields control attentional modulation of alpha and gamma oscillations in contralateral occipitoparietal cortex[J]. Journal of Neuroscience, 2015, 35(4): 1638–1647.

[68] Veniero D, Gross J, Morand S, et al. Top-down control of visual cortex by the frontal eye fields through oscillatory realignment[J]. Nature communications, 2021, 12(1): 1757.

[69] Chang Y-C, Huang S-L. The influence of attention levels on psychophysiological responses[J]. International Journal of Psychophysiology, 2012, 86(1): 39–47.

[70] Sciaraffa N, Borghini G, Di Flumeri G, et al. Joint analysis of eye blinks and brain activity to investigate attentional demand during a visual search task[J]. Brain Sciences, 2021, 11(5): 562.

[71] Sharp P, Melcher D, Hickey C. Endogenous attention modulates the temporal window of integration[J]. Attention, Perception, & Psychophysics, 2018, 80(5): 1214–1228.

[72] Sharp P, Gutteling T, Melcher D, et al. Spatial attention tunes temporal processing in early visual cortex by speeding and slowing alpha oscillations[J]. Journal of Neuroscience, 2022, 42(41): 7824–7832.

[73] García-Pérez MA, Alcalá-Quintana R. On the discrepant results in synchrony judgment and temporal-order judgment tasks: a quantitative model[J]. Psychonomic Bulletin & Review, 2012, 19(5): 820–846.

[74] De Boer-Schellekens L, Eussen M, Vroomen J. Diminished sensitivity of audiovisual temporal order in autism spectrum disorder[J]. Frontiers in integrative neuroscience, 2013, 7: 8.

[75] Kostaki M, Vatakis A. Temporal order and synchrony judgments: a primer for students[G]//Timing and time perception: Procedures, measures, & applications. Brill, 2018: 233–262.

[76] Corbetta M, Patel G, Shulman GL. The reorienting system of the human brain: from environment to theory of mind[J]. Neuron, 2008, 58(3): 306–324.

[77] Cavanagh JF, Frank MJ. Frontal theta as a mechanism for cognitive control[J]. Trends in cognitive sciences, 2014, 18(8): 414–421.

[78] Asanowicz D, Panek B, Kotlewska I, et al. On the relevance of posterior and midfrontal theta activity for visuospatial attention[J]. Journal of Cognitive Neuroscience, 2023, 35(12): 1972– 2001.

[79] Friese U, Daume J, Göschl F, et al. Oscillatory brain activity during multisensory attention reflects activation, disinhibition, and cognitive control[J]. Scientific reports, 2016, 6(1): 32775.

[80] Keller AS, Payne L, Sekuler R. Characterizing the roles of alpha and theta oscillations in multisensory attention[J]. Neuropsychologia, 2017, 99: 48–63.

[81] Murray A, Zerroug Y, Soulières I, et al. The Role of Fronto-Central Theta Oscillations in Inter-Sensory Selective Attention[J]. Psychophysiology, 2025, 62(4): e70055.

[82] Corbetta M, Shulman GL. Control of goal-directed and stimulus-driven attention in the brain[J]. Nature reviews neuroscience, 2002, 3(3): 201–215.

[83] Vossel S, Geng JJ, Fink GR. Dorsal and Ventral Attention Systems: Distinct Neural Circuits but Collaborative Roles[J]. The Neuroscientist, 2014, 20(2): 150–159.

[84] Tamber-Rosenau BJ, Asplund CL, Marois R. Functional dissociation of the inferior frontal junction from the dorsal attention network in top-down attentional control[J]. Journal of Neurophysiology, 2018, 120(5): 2498–2512.

[85] Meyyappan S, Rajan A, Mangun GR, et al. Role of inferior frontal junction (IFJ) in the control of feature versus spatial attention[J]. Journal of Neuroscience, 2021, 41(38): 8065–8074.

[86] Di Dona G, Ronconi L. Beta oscillations in vision: a (preconscious) neural mechanism for the dorsal visual stream?[J]. Frontiers in Psychology, 2023, 14: 1296483.

[87] Engel AK, Fries P. Beta-band oscillations—signalling the status quo?[J]. Current opinion in neurobiology, 2010, 20(2): 156–165.

[88] Fries P. Rhythms for cognition: communication through coherence[J]. Neuron, 2015, 88(1): 220–235.

[89] Palva S, Palva JM. Roles of brain criticality and multiscale oscillations in temporal predictions for sensorimotor processing[J]. Trends in neurosciences, 2018, 41(10): 729–743.

