## Supplemental figure for "Visuospatial attention exerts opposite modulatory effects on explicit and implicit audiovisual subjective synchrony via the frontoparietal network"

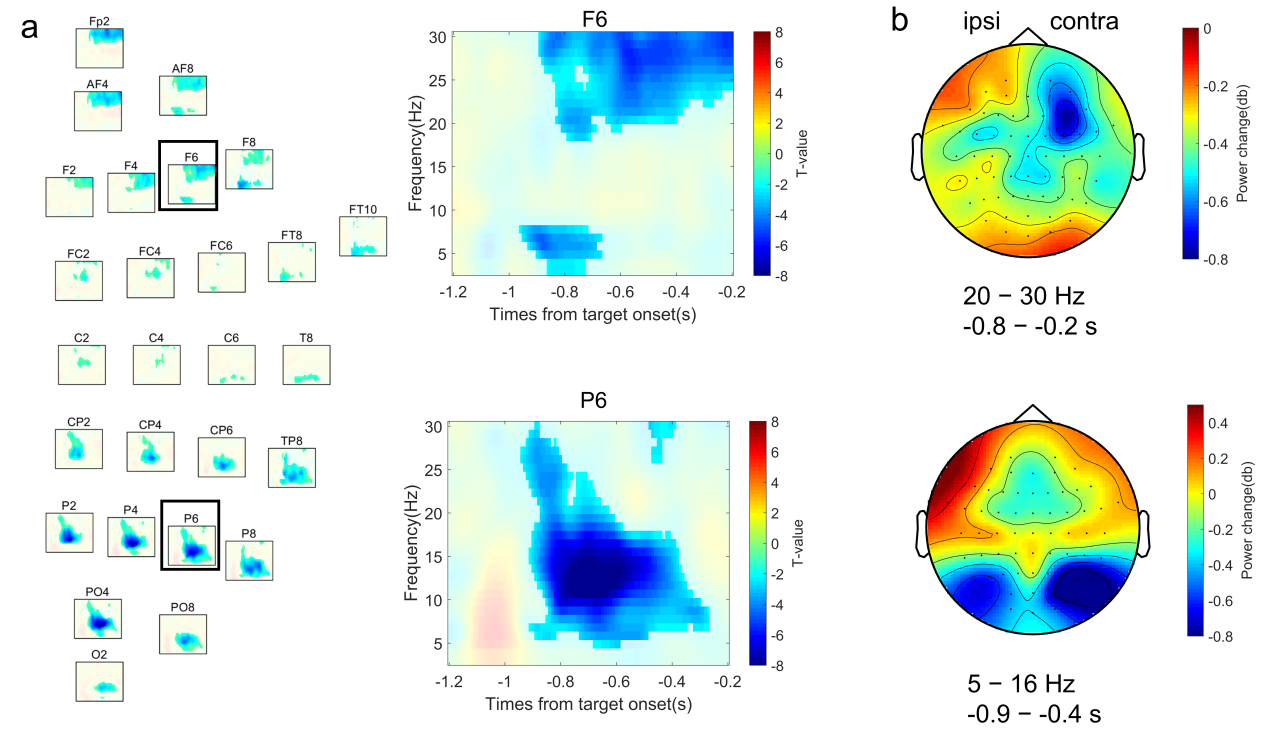

**Fig. S1. Attention-Related Oscillatory Power Changes in Contralateral vs. Ipsilateral Electrodes in Experiment 1.** (a) Left subplot: Time–frequency representation of the paired-sample t-test comparing post-cue (pre-target) oscillatory power between contralateral and ipsilateral electrodes. Unmasked regions indicate significant time–frequency clusters; masked areas represent non-significant results. Electrodes highlighted in the black rectangular frame indicate representative sites. Right subplot: Time–frequency t-value maps for representative electrodes F6 (frontal) and P6 (parietal). (b) Top subplot: Topographical distribution of raw oscillatory power within the 20–30 Hz and −0.8 to −0.2 s time window, revealing marked asymmetry over frontal and frontocentral areas. Bottom subplot: Topographical distribution of raw power in the 5–16 Hz and −0.9 to −0.4 s time window, showing asymmetric activation over centroparietal and parieto-occipital electrodes.

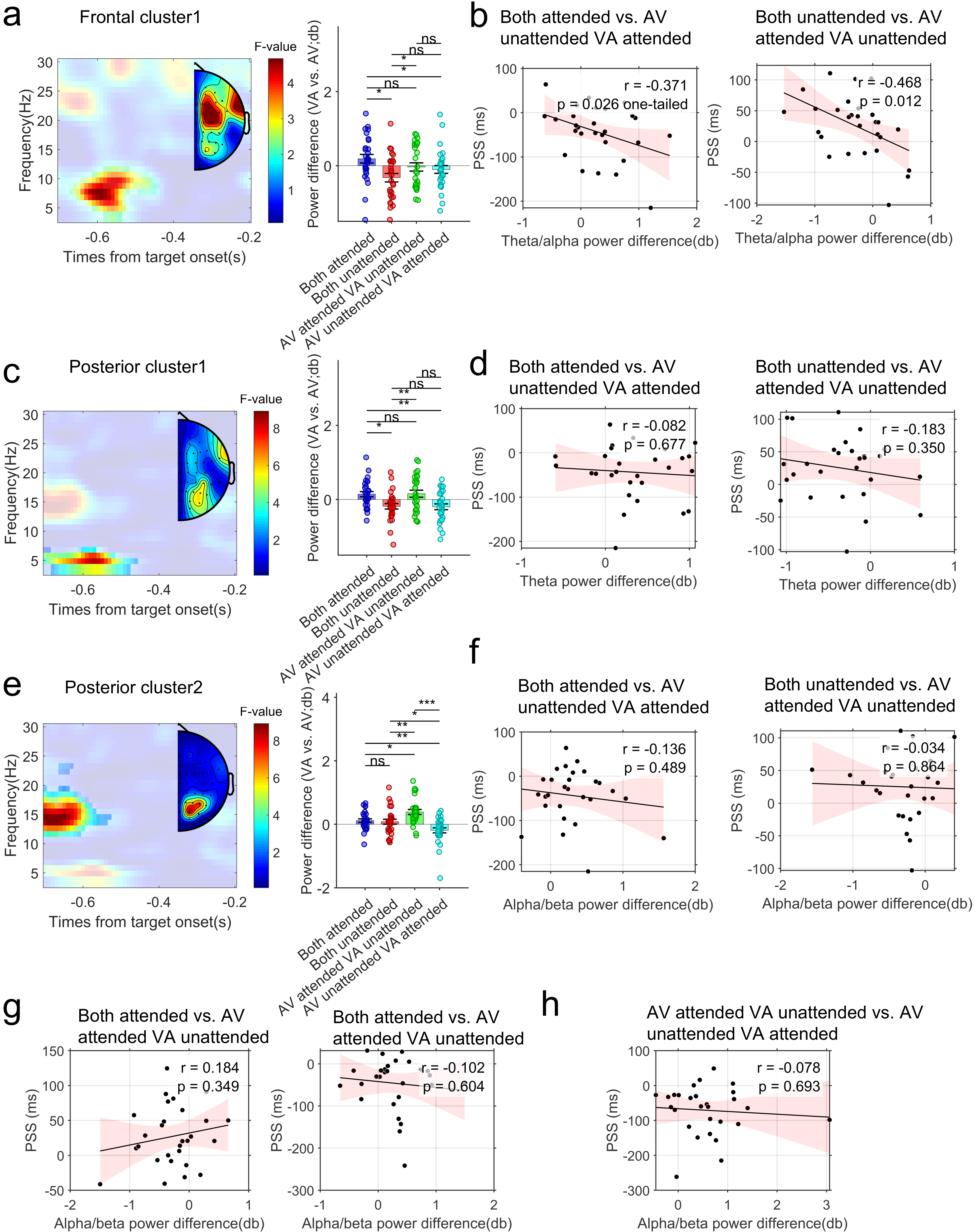

**Fig. S2. Prestimulus Oscillatory Power Differences Across Attentional Configurations and Their Relationship with Behavioral Performance in the the hemispheric analysis in Experiment 1.** (a), (c), and (e) depict the time–frequency representations of the F-statistics for Cluster 1 (frontal[AF4, AF8, F2, F4, F6, FC2, FC4, FC6] and posterior [CP4, CP6, P4, P6, P8] regions) and Cluster 2 (posterior [P2, P4, P6, P8, PO4, PO8] region), along with the post hoc analysis results for the time–frequency windows showing significant effects. Unmasked regions indicate significant time–frequency clusters; masked areas represent non-significant results. (b), (d), and (f–h) display the correlation results between the power differences and PSS differences across attentional configurations that showed significant effects in the post hoc comparisons in panels (a), (c), and (e), respectively. **p* < 0.05,***p* < 0.01,****p* < 0.001. ns indicates not significant.

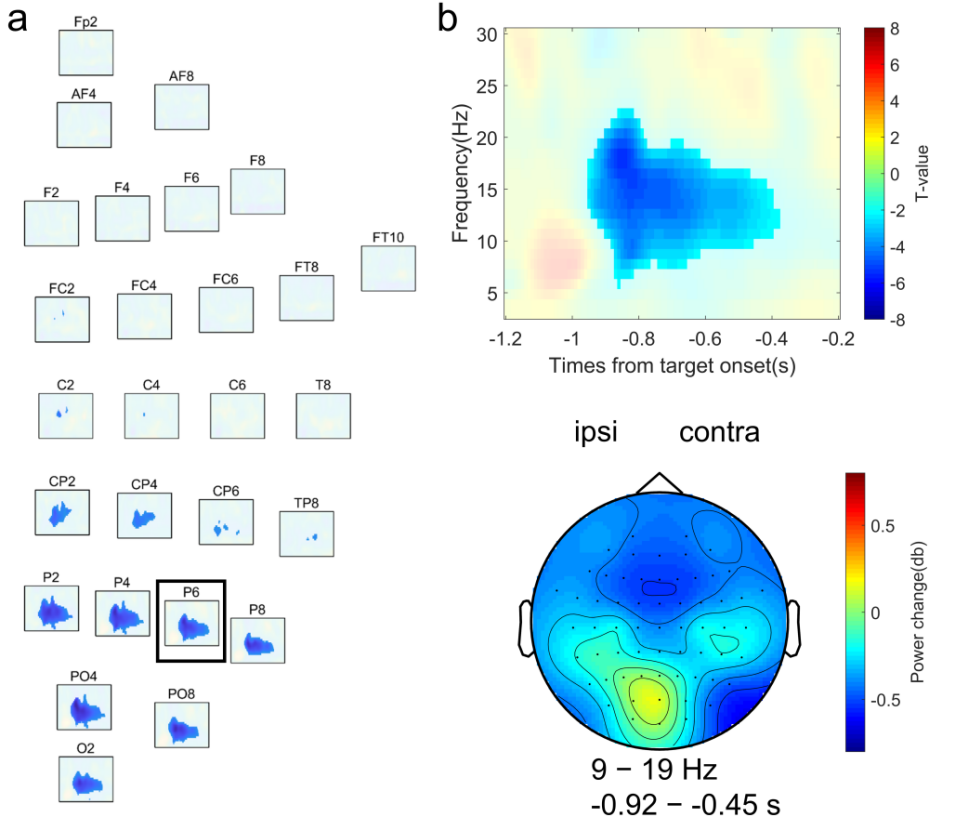

**Fig. S3. Attention-Related Oscillatory Power Changes in Contralateral vs. Ipsilateral Electrodes in Experiment 2.** (a) Time–frequency representation of the paired-sample t-test comparing post-cue oscillatory power between contralateral and ipsilateral electrodes. Unmasked regions indicate significant time–frequency clusters; masked areas represent non-significant results. Electrodes highlighted in the black rectangular frame indicate representative sites. (b) Top subplot: Time–frequency t-value maps for representative electrodes P6 (parietal). Bottom subplot: Topographical distribution of raw power in the 9–19 Hz range and −0.92 to −0.45 s window, showing asymmetric activation over centroparietal and parieto-occipital electrodes.

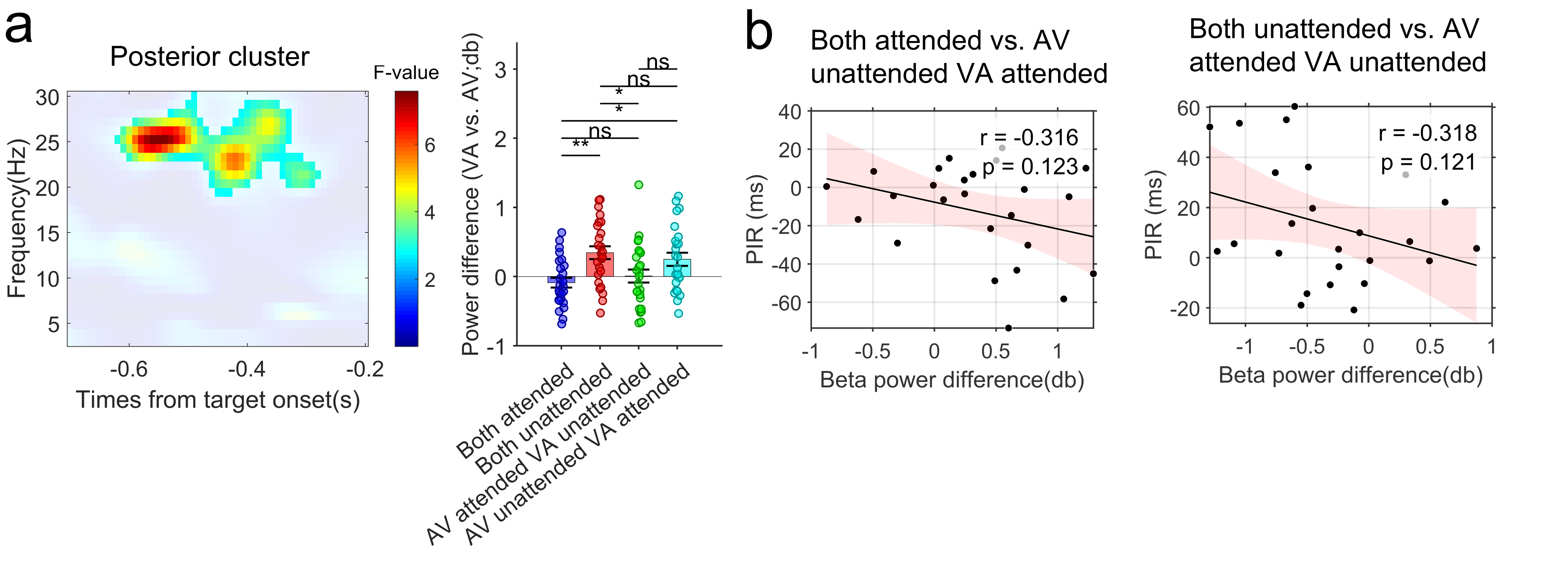

**Fig. S4. Prestimulus Oscillatory Power Differences Across Attentional Configurations and Their Relationship with Behavioral Performance in the the hemispheric analysis in Experiment 2.** (a) The time–frequency representations of the F-statistics in posterior region channels (CP6, CP4, CP6, TP8, P4, P8, PO4, PO8, P6, P2) along with the post hoc analysis results for the time–frequency windows (-0.6 to -0.5 s and 23 to 26 Hz) showing significant effects. Unmasked regions indicate significant time–frequency clusters; masked areas represent non-significant results. (b) The correlation results between the power differences and PSS differences across attentional configurations that showed significant effects in the post hoc comparisons in panels (a) and in PIR analysis. * *p* < 0.05,***p* < 0.01. ns indicates not significant.
